# Parallel multimodal circuits control an innate foraging behavior

**DOI:** 10.1101/326405

**Authors:** Alejandro López-Cruz, Navin Pokala, Aylesse Sordillo, Steven W. Flavell, Patrick T. McGrath, Cornelia I. Bargmann

**Affiliations:** Lulu and Anthony Wang Laboratory of Neural Circuits and Behavior, The Rockefeller University, New York, NY 10065, USA; Department of Biology, Georgia Institute of Technology, Atlanta, GA 30332, USA; Chan Zuckerberg Initiative, Palo Alto, CA 94301, USA

## Abstract

Foraging strategies that enable animals to locate food efficiently are composed of highly conserved behavioral states with characteristic features. Here, we identify parallel multimodal circuit modules that control an innate foraging state -- local search behavior -- after food removal in the nematode *Caenorhabditis elegans*. Two parallel groups of chemosensory and mechanosensory glutamatergic neurons that detect food-related cues trigger local search by inhibiting separate integrating neurons through a metabotropic glutamate receptor, MGL-1. The chemosensory and mechanosensory modules are separate and redundant, as glutamate release from either can drive the full behavior. Spontaneous activity in the chemosensory module encodes information about the time since the last food encounter and correlates with the foraging behavior. In addition, the ability of the sensory modules to control local search is gated by the internal nutritional state of the animal. This multimodal circuit configuration provides robust control of an innate adaptive behavior.

## INTRODUCTION

The optimal foraging strategy across conditions and species can be formulated in a unified framework combining resource distribution, recent experience, and current internal state (Humphries et al., 2010; Nathan et al., 2008; Salvador et al., 2014; Weimerskirch et al., 2007). One prominent foraging strategy within this framework is area-restricted or local search, an intensive exploration over several minutes of the region where food resources were last encountered. As time since the last food encounter passes, animals transition to global search strategies to explore distant areas. The local-to-global search foraging pattern has been observed in fish (Papastamatiou et al., 2012), reptiles (Eifler et al., 2012), insects (Bell, 1985; Lihoreau et al., 2016; Nakamuta, 1985), birds (Dias et al., 2009; Weimerskirch et al., 2007), and mammals (Benedix, 1993; Hills et al., 2013). Although the motor programs employed during local search vary greatly between these species --swimming, crawling, flying, walking -- the underlying behavioral state consisting of a sustained period of intensive searching after food encounter is highly conserved. Understanding the molecular and circuit mechanisms underlying this searching state may provide insight into the basis of ancient conserved behaviors.

Two features of local search hint at circuit elements that may be required for search behavior. First, animals execute local search not only after recent food encounters, but also after encounters with sensory cues associated with resources, such as food odors (Bell, 1985; Murata et al., 2017), light levels (Horstick et al., 2017), and pheromones (Nakamuta, 1985). This observation suggests that local search involves a neuronal representation of food-related sensory features. Second, search behavior is persistent and outlasts its trigger (previous food encounter), suggesting that it results from an internally generated short-term food memory. While the external triggers and dynamics of local search have been extensively characterized, the neuronal circuits that represent the external sensory environment and the sites that hold the short-term food memory are not fully understood.

The compact nervous system of *C. elegans*, which consists of exactly 302 neurons, presents an ideal system to explore these questions. *C. elegans* feeds on bacterial food (Brenner, 1974) and many neurons that detect food-related sensory cues have been characterized. For example, the AWC olfactory neurons detect volatile bacterial odors, ASK neurons detects amino acids (Bargmann, 2006; Gray et al., 2004) and several mechanosensory neurons detect food textures (Albeg et al., 2011; Park et al., 2008; Sawin et al., 2000; Yemini et al., 2013). The synaptic connections between these sensory neurons and downstream interneurons and motor neurons that generate behaviors are known (White et al., 1986), making this an excellent system for defining neuronal circuits related to foraging.

Upon removal from food, *C. elegans* executes a stereotypical local search behavior in which it explores a small area by executing many random, undirected reversals and turns for about fifteen minutes. If it fails to find food, it transitions to a global search in which it explores larger areas by suppressing reorientations and executing long forward runs (Ahmadi and Roy, 2016; Calhoun et al., 2014; Gray et al., 2005; Hills et al., 2004; Salvador et al., 2014; Wakabayashi et al., 2004). The reorientations associated with local search in *C. elegans* are present in different patterns in other behaviors such as chemotaxis (Pierce-Shimomura et al., 1999) and aerotaxis (Hums et al., 2016). Previous studies have identified sensory neurons that contribute to acute reorientations during local search by activating glutamate-gated ion channels on downstream neurons (Chalasani et al., 2007; Gray et al., 2005; Hills et al., 2004; Wakabayashi et al., 2004). However, ion channels signal over sub-second timescales, whereas local search occurs over minutes. The relationship between acute sensory signals and the longer-term dynamics of the local search state is not known.

Here, we identify a G protein-coupled glutamate receptor, MGL-1, as a transducer of sensory information into a sustained circuit state. We show that parallel chemosensory and mechanosensory modules have redundant capability to generate the local search behavioral state, and that each module signals food removal by inhibiting separate downstream integrating neurons through MGL-1. The chemosensory module encodes time since food removal in its spontaneous activity patterns, but drives local search only in the context of a third signal representing an internal satiety state. Our findings reveal a circuit organization that allows for the robust execution of a conserved adaptive behavior.

## RESULTS

### Local search behavior in wildtype *C. elegans*

We studied local search behavior by transferring animals from a standard bacterial food lawn to a large agar plate without food (~80 cm^2^), where we filmed and quantified their behavior continuously for 45 min (Fig. 1A). As described previously (Calhoun et al., 2014; Gray et al., 2005; Hills et al., 2004; Wakabayashi et al., 2004), animals initially explored a small area by performing many reorientations (Fig. 1B-C, local search), then gradually transitioned to exploring a larger area by performing fewer reorientations over time (Fig. 1B-C, global search). *C. elegans* can perform a variety of reorientations using different locomotor sequences (Fig. S1). We found that reorientations that start with a reversal were increased during local search (time-dependent, Fig S1A-C), but reorientations resulting from turning during forward movement occurred at similar frequencies during local and global search (time-independent, Fig. S1D-E). Here, we focused on the time-dependent reorientations that drive local search behavior (Fig. 1C).

**Figure 1.**
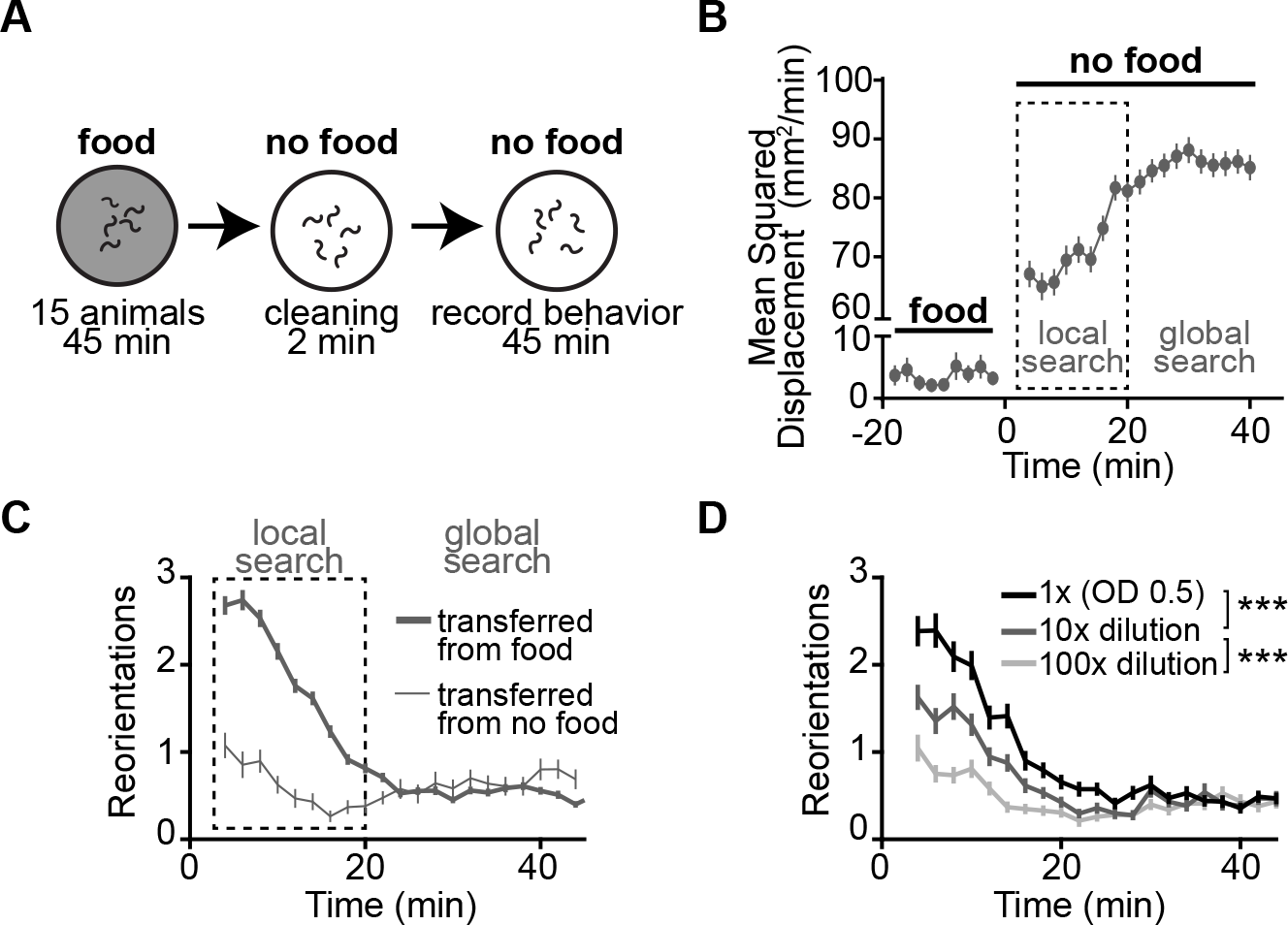
Off-food foraging in wildtype animals. (A) Schematic of off-food foraging assay. Animals were transferred from a homogeneous bacterial food lawn to a large plate (80 cm^2^) without food, where their behavior was filmed for 45 min. The behavior varies from day to day, so all experiments were compared to control animals recorded simultaneously in an adjacent behavioral chamber. (B) Average mean squared displacement per minute on food and after food removal. (C) Thick line: off-food foraging showing mean reorientations in 2-min time windows after food removal. The first two minutes are not plotted as these are affected by transfer (Zhao et al., 2003). Thin line: similar animals transferred after 45 min on a plate without food. (D) Effect of food concentration on subsequent off-food foraging. Animals were removed from either a standard OD=0.5 bacterial lawn (n=103), a lawn diluted 10-fold (n=114), or a lawn diluted 100-fold (n=100), and their off-food behavior was recorded. All data are presented as mean ± s.e.m. ***p < 0.001 by Wilcoxon rank sum test with Bonferroni correction.

If local search represents a short-term memory of recent food experience, it should be modulated by food history. To test this prediction, we varied the food concentration that animals experienced prior to food removal. Animals that had been in dilute food performed fewer reorientations during local search than animals that had been in concentrated food (Fig. 1D). Reorientations during global search remained unchanged for all initial conditions. This result and others (Calhoun et al., 2015), suggest that local search is a memory state that is actively generated and modulated in response to food removal, and that it transitions or decays over time to a memoryless, default global search state.

The transition from local to global search is gradual at a population level (Fig. 1C). To characterize the transition in individuals, we looked at reorientation dynamics for 1,631 single animals and searched for abrupt transitions in reorientation frequency (Fig. S2A-C). 51% of animals exhibited a single abrupt transition corresponding to the end of local search (Fig. S2A-C), whereas 49% exhibited multiple transitions or no clear transition (Fig. S2A-B). There was a wide range in the duration of local search and in the intensity of local search in individual animals (Fig. S2C-D). Despite this variability, 93% of the animals performed more reorientations at early times after food removal (Fig. S2E), indicating that local search is a robust behavioral response to a recent food encounter.

### The metabotropic glutamate receptor, *mgl-1*, is necessary for local search

To gain insight into the neuronal circuit mechanisms that generate local search behavior, we tested 42 candidate mutant strains lacking individual ionotropic and metabotropic neurotransmitter receptors by filming their behavior and comparing their reorientation rates to matched controls. For each mutant strain, we calculated the average fold-change in reorientations relative to wildtype during local search (2-20 min) (Fig. 2A, Table S1) and global search (30-45 min) (Table S1). We focused on mutants that had stronger defects during local search. Of the 42 mutant strains tested, two notable findings emerged: a set of ionotropic acetylcholine receptor mutants (*lgc-46, acc-4, acr-12*, Fig. S3A, Table S1), and the mutant strain CX17083 (Fig. 2B, Table S1).

**Figure 2.**
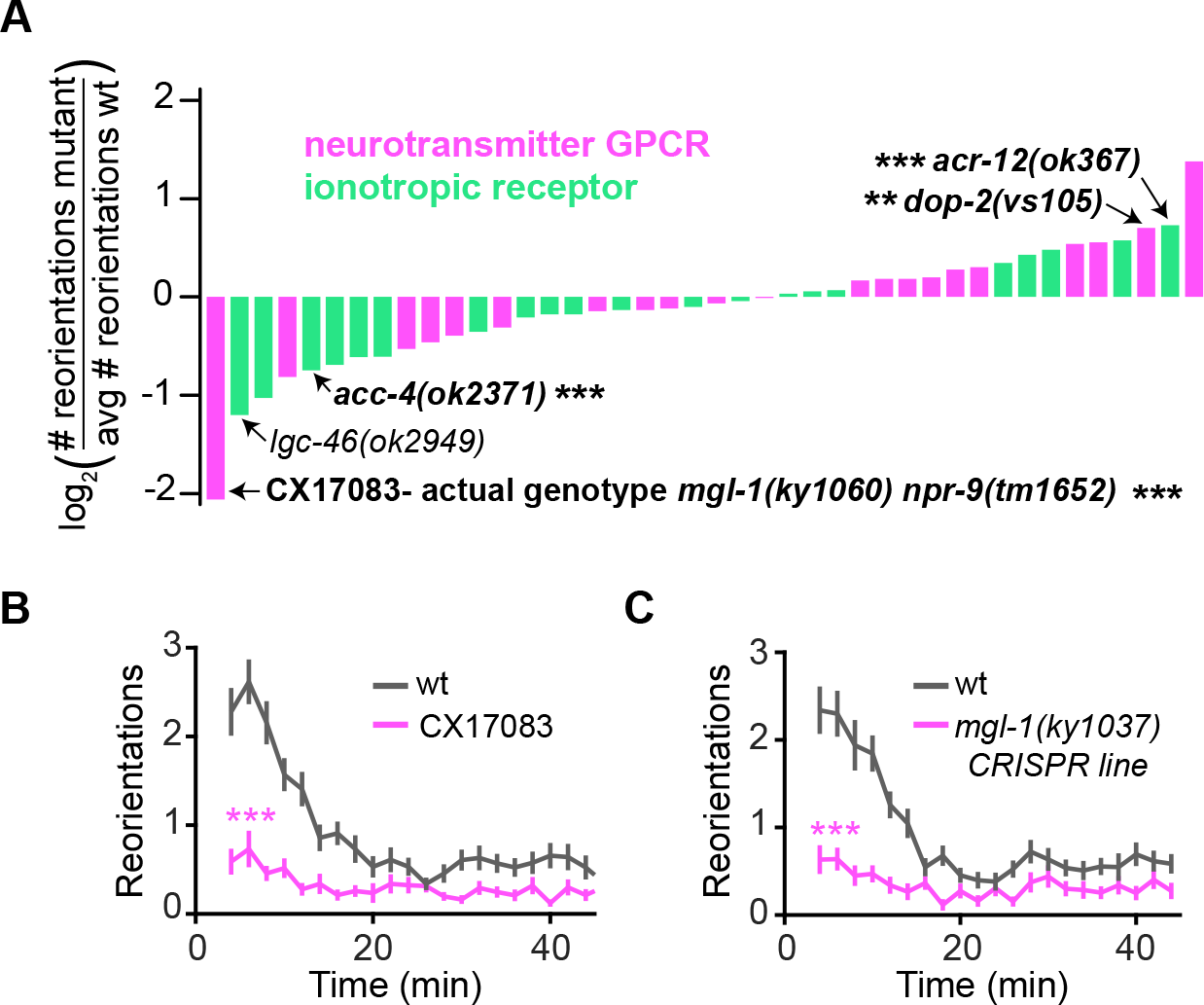
*mgl-1*, a metabotropic glutamate receptor, is necessary for local search. (A) Local search (2-20 min) reorientation rates normalized to wildtype controls for 42 mutant strains. Data shows the fold change in reorientations during local search for each mutant strain, presented as the log2 of mutant/wildtype reorientation ratio. Mutants bolded and marked with the p-value asterisks met two criteria: (i) showed a statistically significant fold change during local search of at least 1.5 (abs(log_2_(ratio))>0.58), and (ii) local search was affected more than global search. For detailed results and explanations, see Table S1. (B) Off-food foraging for mutant strain CX17083 [*npr-9(tm1652);mgl-1(ky1060)*] (wt, n =56; CX17083, n=62). Presented as mean ± s.e.m. ***p < 0.001 by Wilcoxon rank sum test. (C) Off-food foraging for CRISPR-generated *mgl-1(ky1037)* (wt, n=61; *mgl-1(ky1037),* n=41). Presented as mean ± s.e.m. ***p < 0.001 by Wilcoxon rank sum test.

Mutants lacking the inhibitory acetylcholine receptors *lgc-46* and *acc-4* showed diminished local search (Fig. S3A), whereas mutants in the excitatory acetylcholine receptor *acr-12* showed augmented local search behavior (Fig. S3A). These results suggest an involvement of acetylcholine in controlling reorientation probabilities during local search. *lgc-46* and *acc-4* have been hypothesized to form heteromeric channels and are both expressed broadly in cholinergic neurons (Pereira et al., 2015; Takayanagi-Kiya et al., 2016). We were unable to rescue *lgc-46* in specific subset of neurons (cholinergic motor neurons, sets of interneurons, data not shown) to uncover a circuit mechanism, and therefore did not study the cholinergic mutants further.

CX17083 was profoundly defective in local search, with milder defects in reorientation rates during global search (Fig. 2B, Table S1). It was originally selected for its annotated *npr-9(tm1552)* mutation (Bendena et al., 2008) but we were unable to rescue its local search defect by rescuing functional *npr-9* expression with a transgene (Fig. S3B), and a new CRISPR-generated *npr-9* mutant allele did not recapitulate the phenotype (Fig. S3C), suggesting that a background mutation caused the observed phenotype. To identify the unmapped background mutation, we sequenced CX17083 and found a large deletion within *mgl-1,* a G-protein coupled inhibitory metabotropic glutamate receptor (Fig. S3D) (Dillon et al., 2006). Restoring *mgl-1* expression by injecting a *mgl-1* transgene into CX17083 rescued the mutant behavior (Fig. S3E), and a new CRISPR-generated *mgl-1* mutant allele recapitulated the CX17083 phenotype (Fig. 2C), confirming that *mgl-1* is the causative mutation affecting local search. Our results indicate that *mgl-1* is necessary to generate local search. *mgl-1* has been implicated in food-related physiological responses (Greer et al., 2008; Kang and Avery, 2009a, b), pharyngeal pumping (Dillon et al., 2015), and foraging (Ahmadi and Roy, 2016), making it a good entry point to circuit mechanisms related to food-related behaviors.

### *mgl-1*inhibits AIA and ADE to generate local search

To identify the circuits where *mgl-1* acts to generate local search, we rescued *mgl-1* in subsets of its endogenous expression pattern using an intersectional transgene approach (Fig. 3A). A transgene containing an inverted, inactive floxed *mgl-1* genomic fragment was activated in a subset of cells by a second transgene expressing Cre recombinase under cell-specific promoters. Successful expression was confirmed by GFP (Fig. 3A). Activating the *mgl-1* fragment in all neurons rescued the *mgl-1* local search defect (Fig. 3B) and showed broad *mgl-1* GFP expression. To narrow down the sites of *mgl-1* action, we activated the *mgl-1* fragment in a smaller subset of cells using a *mod-1::nCre* transgene (Fig. S4A). Simultaneous activation in sensory neurons ASI, Il1, and ADE, the interneuron AIA, and the motor neurons RMD and NSM rescued local search (Fig. S4A). Of these neurons, activating *mgl-1* in AIA and ADE or both neurons (Fig. 3C), but not in RMD, ASI, NSM, or IL1 (Fig. S4B), was sufficient for rescue. In control experiments, transgenes with nonsense mutations in the *mgl-1* coding region failed to rescue the behavior (Fig. S4C-D). *mgl-1* had been previously reported to be expressed in AIA, but not in ADE. The expanded expression we observe is likely due to the use of a genomic fragment that includes all introns.

**Figure 3.**
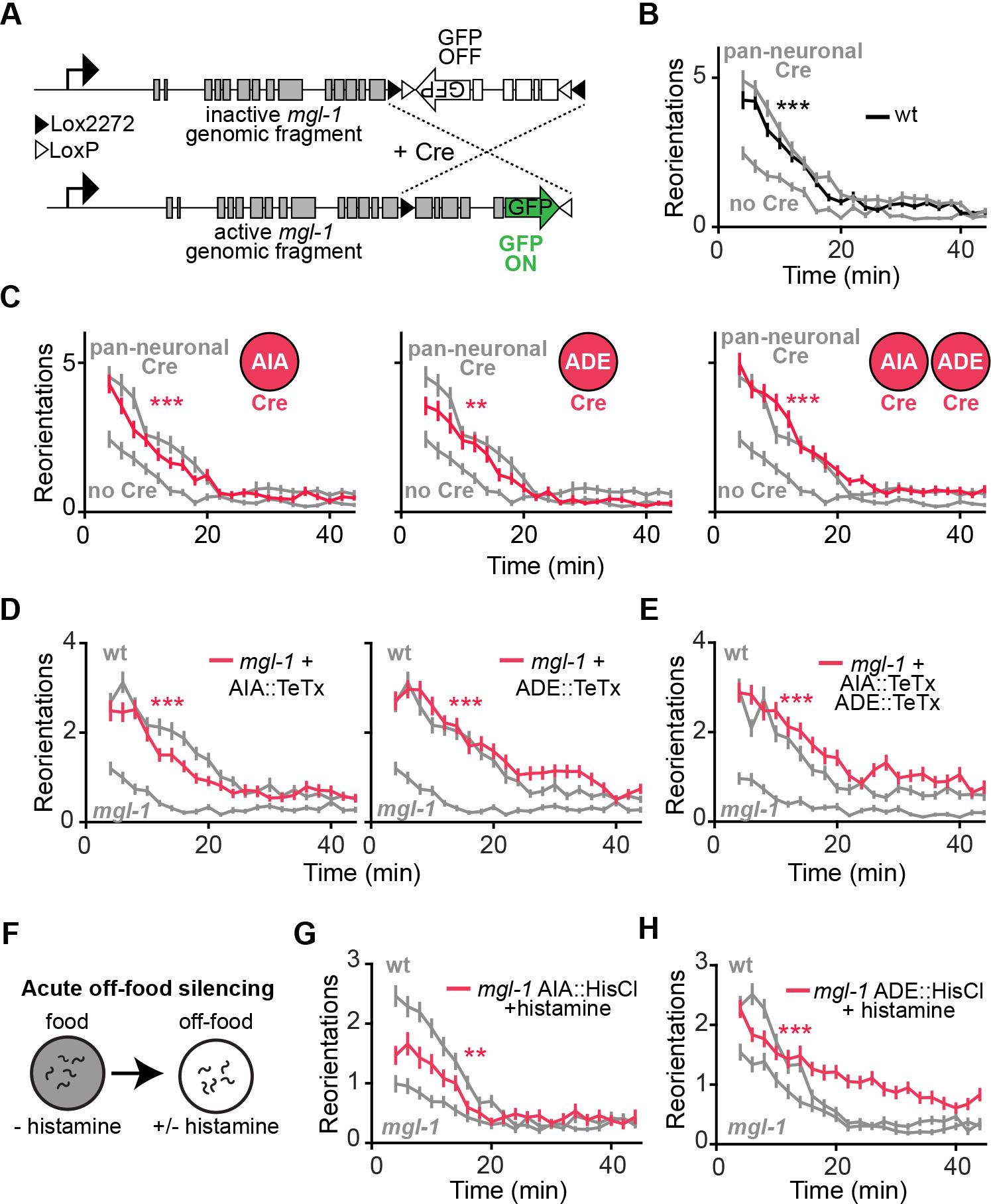
*mgl-1* silences AIA and ADE to generate local search. (A-C) *mgl-1* rescue in subsets of its endogenous expression pattern using an intersectional transgene approach. (A) Schematic depicting Cre-Lox transgene approach. Successful activation is confirmed by GFP, which is expressed with *mgl-1* in a bicistronic transcript. (B) Rescue of *mgl-1* by reconstituting inverted transgene in all neurons using pan-neuronal Cre (*tag-168::nCre*). Animals that only express the inverted transgene are labelled ‘no Cre’ (no Cre, n=72; pan neuronal Cre, n=69). (C) Rescue of *mgl-1* by expression in AIA, ADE or both neurons (no Cre, n= 46; AIA rescue, n=50; ADE rescue, n=32; AIA+ADE rescue, n=48). (D) Rescue of *mgl-1(ky1037)* phenotype by expression of tetanus toxin light chain (TeTx) in AIA or ADE. (*mgl-1*, n=78; AIA::TeTx *mgl-1*, n = 63; ADE::TeTx *mgl-1*, n=82). (E) Rescue of *mgl-1(ky1037)* phenotype by expression of tetanus toxin light chain (TeTx) in both AIA and ADE (*mgl-1*, n=72; AIA::TeTx+ ADE::TeTx *mgl-1*, n=69). (F) Acute neuronal silencing by expressing cell-specific histamine-gated chloride channel and adding exogenous histamine to plates. Negative controls are *mgl-1(ky1037)* animals expressing the histamine gated chloride channel that were transferred to plates without histamine. (G) Acutely silencing AIA off-food gives partial rescue of *mgl-1(ky1037)* phenotype (*mgl-1* AIA::HisCl-his, n=84; *mgl-1* AIA::HisCl +his, n=89). (H) Acutely silencing ADE and other dopaminergic neurons (CEP, PDE) off-food gives partial rescue of local search and an altered pattern of global search behavior. (*mgl-1* ADE::HisCl-his, n=113; *mgl-1* ADE::HisCl +his, n=111). The dopaminergic neurons are known to affect locomotion speed and turning. All data presented as mean ± s.e.m. p-values calculated using Wilcoxon rank sum test with Bonferroni correction. **p < 0.01 ***p < 0.001.

As *mgl-1* is predicted to be an inhibitory metabotropic receptor, animals that lack *mgl-1* may have defective local search because they fail to silence the AIA and ADE neurons. To test this possibility, we expressed tetanus toxin light chain in AIA, ADE, or both neurons in *mgl-1* mutant animals (Fig. 3D-E). Inhibiting neurotransmitter release from either AIA, ADE, or from both neurons with tetanus toxin rescued local search behavior in *mgl-1* (Fig. 3D-E). Tetanus toxin expression in another *mgl-1*-expressing neuron, NSM, did not rescue local search (Fig. S4E).

Neurons expressing tetanus toxin lack synaptic vesicle release chronically throughout life. To silence AIA or ADE acutely during foraging behavior, we expressed the *Drosophila* histamine-gated chloride channel HisCl1 in either neuron (Pokala et al., 2014). *C. elegans* does not use histamine as an endogenous transmitter, so selective expression of HisCl1 permits rapid silencing of target neurons by adding histamine to the assay plate (Fig. 3F). Acutely silencing AIA off food partially rescued the local search defect in *mgl-1* mutants, albeit to a lesser extent than tetanus toxin (Fig. 3G). There are no ADE-specific promoters; acutely silencing ADE plus other dopaminergic neurons partly rescued local search and also increased reorientation frequencies at later time points after food removal, leading to altered global search (Fig. 3H).

Together these results indicate that when animals are removed from food, inhibition of AIA or ADE via MGL-1 generates local search. This interpretation in turn suggests that AIA and ADE normally suppress reorientations, and indeed an inhibitory effect of AIA on reorientations had been previously reported (Wakabayashi et al., 2004). AIA and ADE each release multiple neurotransmitters and peptides that could mediate this behavioral effect. We tested two candidates, dopamine from ADE and *ins-1* from AIA, but found that neither explains local search behavior (Fig. S4F) (see Discussion).

### Parallel, multimodal glutamatergic sensory pathways can redundantly generate foraging behavior

We next focused on the sources of glutamate that silence AIA and ADE through MGL-1 to generate local search. AIA and ADE do not express *eat-4*, the vesicular glutamate transporter that loads glutamate into synaptic vesicles for release (Serrano-Saiz et al., 2013), but they are postsynaptic to multiple glutamatergic neurons. The wiring diagram of *C. elegans* (White et al., 1986) and *eat-4* expression data (Serrano-Saiz et al., 2013) predict that AIA receives glutamatergic input from several chemosensory neurons that detect food-related volatile odors and soluble chemicals (Chalasani et al., 2007; Wakabayashi et al., 2009) (Fig. 7). ADE is predicted to receive glutamatergic input from mechanosensory neurons regulated by food texture (Albeg et al., 2011; Serrano-Saiz et al., 2013; White et al., 1986; Yemini et al., 2013) (Fig. 7).

To characterize the role of sensory glutamate inputs in local search behavior, we modified the endogenous *eat-4* (VGLUT1) locus of wild-type animals to permit cell-specific knockout of glutamate release. Two successive rounds of CRISPR/Cas9 editing were performed to insert an FRT before the start codon and *let-858*-3̍-UTR::FRT::mCherry after the stop codon of the endogenous *eat-4* coding region (Fig. 4A) (Schwartz and Jorgensen, 2016). In this edited *eat-4* strain, expression of a nuclear localized flippase (nFLP) under cell-specific promoters excised the *eat-4* coding region and resulted in the appearance of mCherry in the targeted cells (Fig. 4A). To validate this approach, we expressed nFLP under a pan-neuronal promoter (*tag-168::nFLP*); this resulted in a strong reorientation defect reminiscent of *eat-4* mutants (Hills et al., 2004) (Fig. 4B) and neuronal mCherry expression consistent with *eat-4* expression patterns (Fig. S5A-B).

**Figure 4.**
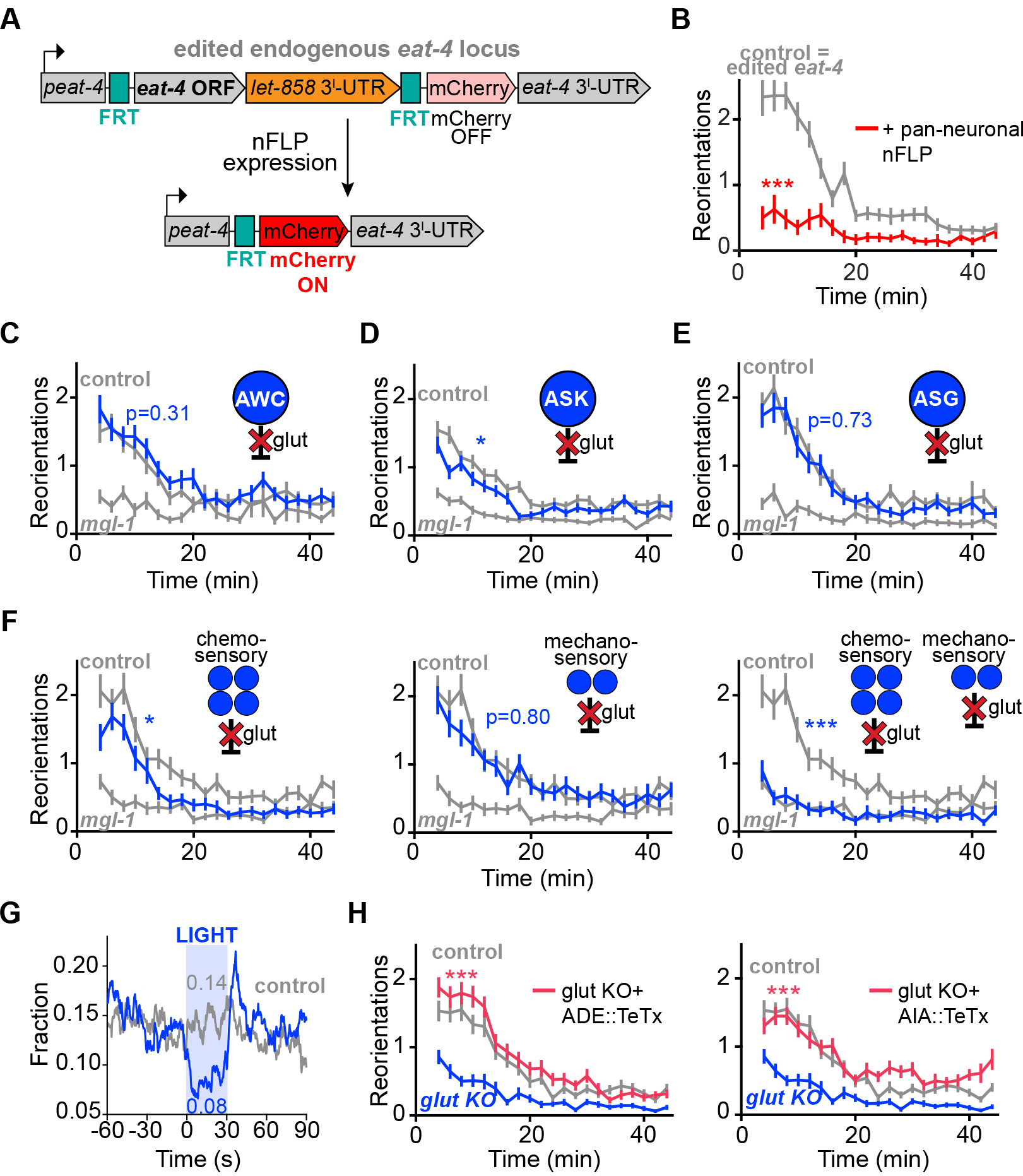
Parallel, multimodal glutamatergic sensory pathways redundantly generate local search. (A) Schematic of endogenous glutamate knockout strategy. Using CRISPR/Cas9, an FRT site was inserted immediately before the start codon of *eat-4* (VGLUT1), and *let-858* 3̍-UTR:: FRT::mCherry immediately after the stop codon of *eat-4*. *let-858* 3̍-UTR stops transcription so mCherry is not expressed. To knock out glutamate release in this edited strain, nls-flippase (nFLP) was expressed under cell-specific promoters, leading to excision of the *eat*-*4* ORF, confirmed by mCherry expression in the targeted cells. (B) Validation of glutamate knockout strategy. Pan-neuronal glutamate knock out by expressing pan-neuronal nFLP (*tag-168::nls-FLP)* in the edited *eat-4* strain. Animals show phenotype reminiscent of *eat-4* mutants (control, n=65; pan-neuronal nFLP, n=56). (C-E) Off-food foraging after glutamate knockout from individual chemosensory neurons. (C) AWC glutamate knockout (control, n=55; AWC glut KO, n=57). (D) ASK glutamate knockout (control, n=108; ASK glut KO, n=92). (E) ASG glutamate knockout (control, n=60; ASK glut KO, n=58). (F) Off-food foraging after glutamate knockout from all chemosensory neurons, all mechanosensory neurons, or both. (control, n=59; chemo glut KO, n=65; mechano glut KO, n=60; chemo+mechano glut KO, n=53). (G) Light-suppressed reorientations during local search in animals expressing the light-activated chloride channel, *Gt*ACR2 in glutamatergic chemosensory and mechanosensory neurons that synapse onto AIA and ADE (AWC, ASK, ASE, ASG, FLP, AVM). Controls express *Gt*ACR2 and are treated with light, but not pre-incubated with retinal. Data shows the instantaneous fraction of animals performing a reorientation. The mean fraction during stimulation is reported on the plot. Light stimulations were delivered during local search (2-14 min after food removal). (sensory *Gt*ACR2, n=151; controls, n=159). (H) Off-food foraging for both mechano- and chemo- sensory glutamate knockout (glut KO), plus ADE (left) or AIA (right) inhibition with TeTx. (chemo+mechano glut KO, n=77; chemo+mechano glut KO + ADE::TeTx, n= 107; chemo+mechano glut KO + AIA::TeTx, n=77). All data except (G) presented as mean ± s.e.m. p-values calculated using Wilcoxon rank sum test with Bonferroni correction. *p<0.05 ***p < 0.001.

We began by knocking out glutamate release from the individual chemosensory neurons AWC, ASK and ASG, which respond to the removal of chemosensory cues related to food (Chalasani et al., 2007; Wakabayashi et al., 2009). None of these single cell glutamate knockouts resulted in a striking behavioral defect (Fig. 4C-E). Next, we knocked out glutamate release more broadly: from all chemosensory glutamate neurons that synapse onto AIA, from all mechanosensory glutamate neurons that synapse onto ADE, or from both pathways simultaneously. Knocking out glutamate release from either sensory pathway individually had little effect on local search (Fig. 4F, left and middle panels). Only glutamate knockout from both the chemosensory and mechanosensory pathways resulted in a local search defect, which was indistinguishable from that of *mgl-1* mutants (Fig. 4F, right panel, Fig. S5C).

To relate acute activity in these sensory pathways to local search behavior, we transiently silenced the relevant glutamatergic sensory neurons of animals performing local search using optogenetics (Govorunova et al., 2015). We expressed the light-gated chloride channel *Gt*ACR2 in the glutamatergic chemosensory and mechanosensory neurons that synapse onto AIA and ADE (Fig. 7) by using an inverted Cre-Lox recombination strategy (See Methods). Silencing the neurons for 30 sec with light led to a nearly two-fold decrease in reorientation probability, indicating that sensory activity is necessary to maintain a high reorientation state during local search (Fig. 4G).

The local search defect after chemosensory and mechanosensory glutamate knockout might be due to loss of glutamate necessary to inhibit AIA or ADE, or to loss of glutamate signaling to other neurons. To distinguish between these possibilities, we inhibited AIA or ADE with tetanus toxin in animals lacking glutamate in both sensory pathways. Tetanus toxin expression in either AIA or ADE fully restored local search behavior (Fig. 4H), supporting the importance of AIA and ADE as targets of sensory glutamate release. Taken together, these results suggest that glutamatergic chemosensory and mechanosensory neurons can each independently drive full local search behavior by inhibiting either AIA or ADE.

### Spontaneous activity in ASK and AIA reports time after food removal

Acute food removal leads to a transient activation of multiple glutamatergic sensory neurons (Chalasani et al., 2007; Wakabayashi et al., 2009). To investigate the neuronal activity associated with long-term food removal, we monitored spontaneous calcium activity in animals expressing genetically encoded calcium indicators in ASK sensory neurons and AIA interneurons (Akerboom et al., 2012) (Fig. 7). Of the chemosensory neurons, ASK was selected because it was the only one to cause a significant, albeit mild, behavioral defect when *eat-4* was knocked out (Fig. 4D), and because it is directly activated by food removal (Calhoun et al., 2015; Wakabayashi et al., 2009). Animals were removed from food and loaded into a microfluidic device (Chronis et al., 2007), where the spontaneous activities of ASK and AIA were monitored either individually (Fig. 5A-B) or simultaneously (Fig. 5G) for 10 min immediately after food removal (0-10 min, local search) and again after 40 min (40-50 min, global search) (Fig. S6A).

**Figure 5.**
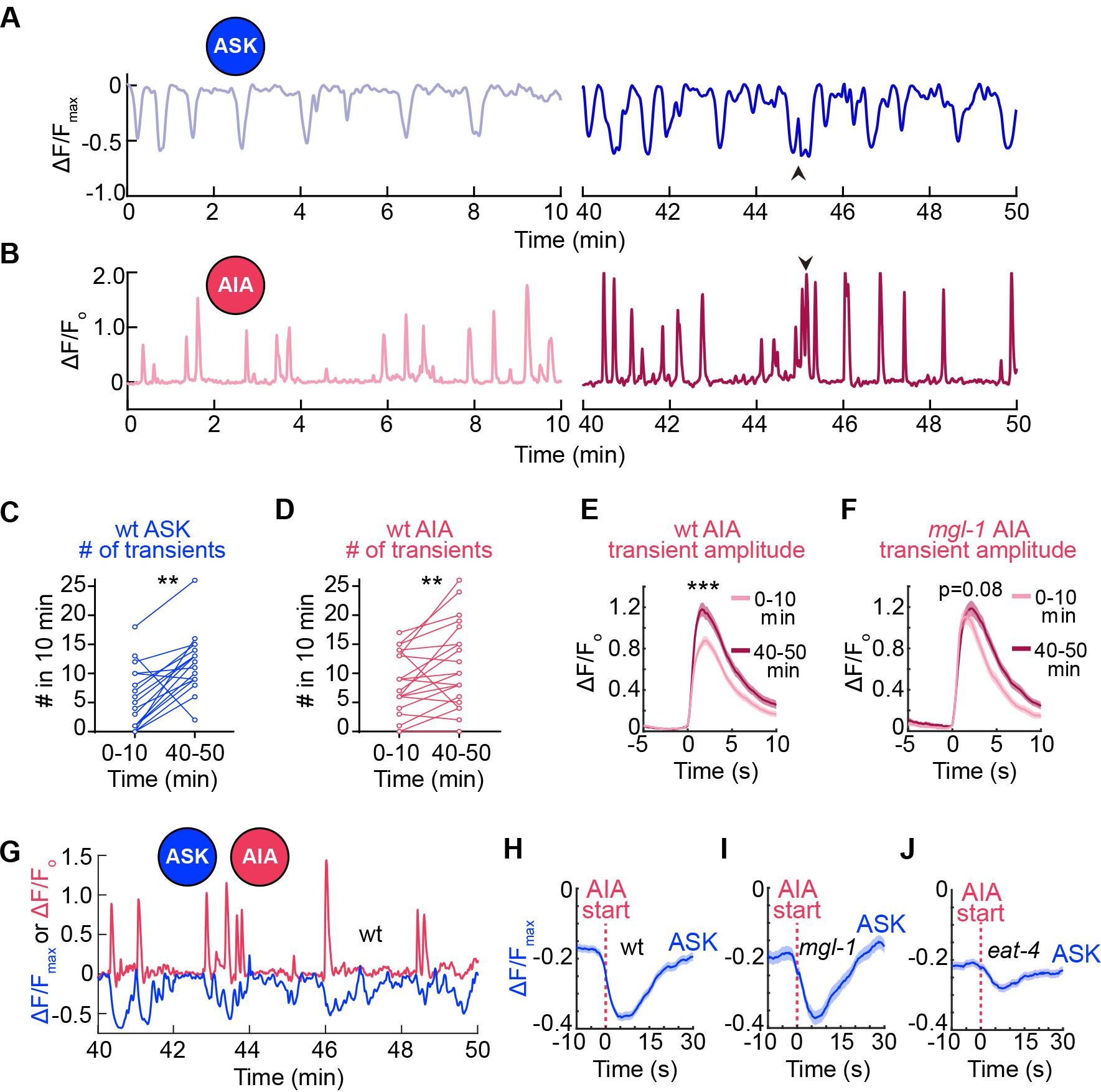
Neuronal dynamics change over time since food removal. (A-B) Spontaneous off-food calcium activity in ASK or AIA neurons. (A) Spontaneous calcium dynamics in a representative ASK neuron (A) or AIA neuron (B) expressing GCaMP5A, recorded 0-10 min and again 40-50 min after food removal. (C) Number of ASK negative transients in each animal early (0-10 min) vs. late (40-50 min) after food removal for wildtype (n=19). **p<0.01 by Wilcoxon signed rank test. Data obtained from imaging ASK individually. (D) Number of AIA positive transients in each animal early (0-10 min) vs. late (40-50 min) after food removal for wildtype. (n=21). **p<0.01 by Wilcoxon signed rank test. Data obtained from imaging AIA individually. (E) Aligned AIA positive transients early (0-10 min, n=210) vs. late (40-50 min, n=255) after food removal for wildtype animals. ***p<0.001 for difference in amplitude by Kolmogorov-Smirnov test. Data obtained from imaging AIA individually. (F) Aligned AIA positive transients early (0-10 min, n=137) vs. late (40-50 min, n=240) after food removal for *mgl-1* animals. p-values for difference in amplitude calculated using Kolmogorov-Smirnov test. Data obtained from imaging AIA individually. (G) Representative trace of single wildtype animal in buffer, showing spontaneous calcium dynamics in simultaneously recorded ASK and AIA neurons expressing GCaMP5A, 40-50 min after food removal. (H-J) ASK traces aligned to the start of AIA transients in wildtype (H) (n=19 animals, 248 transients), *mgl-1* mutants (I) (n=12 animals, 89 transients), and *eat-4* mutants (J) (n=21 animals, 410 transients). Data presented as mean ± s.e.m (shaded region). Data obtained from imaging ASK and AIA simultaneously 40-50 min after food removal.

Both ASK and AIA exhibited unexpected spontaneous activity in the absence of food or obvious external stimuli (Fig. 5A-B). ASK had spontaneous decreases in calcium levels from a high baseline (negative transients) (Fig. 5A), whereas AIA had spontaneous increases in calcium levels from a low baseline (positive transients) (Fig. 5B). Negative transients in ASK lasted between 4-80 sec (median=21.0 sec, Fig. S6B), whereas positive transients in AIA lasted between 3-20 sec (median=7.0 sec, Fig. S6C). The longer transients in both neurons corresponded to complex activity consisting of multiple events (arrowheads, Fig. 5A-B).

To investigate the relationship between these neurons, we imaged their activity simultaneously (Fig. 5G). Spontaneous transients in ASK and AIA often occurred at the same time (Fig. 5G), with ASK negative transients coinciding with the start of AIA positive transient onsets (Fig. 5H). In agreement with this observation, the two neurons showed a strong negative correlation in activity (Fig. S7B).

We next examined the activity of ASK and AIA at the early and late times after food removal associated with local and global search. Spontaneous negative transients in ASK became more frequent at later times after food removal (Fig 5C). This reduction in average ASK activity would be predicted to decrease reversal rates (see optogenetic silencing in Fig. 4G), as is observed in global search. In agreement with its reciprocal activity patterns, AIA had more positive transients at the later times (Fig. 5D). In addition, the AIA transients (but not ASK transients) at late time points were larger in amplitude than early transients (Fig. 5E, Fig. S6D-E). Together, these results indicate that ASK and AIA activity states report information about the time since food removal, an essential component of local search behavior.

### Glutamate shapes activity patterns at multiple time scales

To investigate the role of *mgl-1* and glutamate in the observed spontaneous activity patterns, we imaged ASK and AIA simultaneously in *mgl-1* and *eat-4* mutants. Like wildtype animals, animals lacking *mgl-1* showed anticorrelated activity in ASK and AIA (Fig. 5I, Fig. S7C), and an increased number of AIA positive transients at later times (Fig. S7A, left panel). Thus *mgl-1* is not required for spontaneous AIA activity, for its entrainment by time off food, or for the fast coupling of ASK and AIA. Unlike wildtype, however, AIA transients in *mgl-1* animals had similar amplitudes at early and late time points after food removal (Fig. 5F, Fig. S7A right panel). This result suggests that *mgl-1* may reduce the gain of the ASK-to-AIA synapse during local search.

Like *mgl-1* mutants, mutants in the glutamate transporter EAT-4 did not show a difference in AIA transient amplitude at early and late time points after food removal (Fig. S7E). In addition, *eat-4* animals lacking glutamate release showed decorrelation of ASK and AIA activity (Fig. 5J, Fig. S7D). The decreased correlation was not due to differences in the number, size, or duration of *eat-4* ASK transients, as those were indistinguishable from wildtype (Fig. S7G). Instead, this result suggests that additional glutamate receptors other than MGL-1, most likely glutamate-gated ion channels, mediate fast inhibitory synapses from ASK and AIA. The overall magnitude of AIA transients in *eat-4* mutants was also reduced, consistent with a wider role for glutamate signaling at AIA synapses (Fig. S7F). Together these results suggest that glutamate acts through multiple receptors including *mgl-1* to modulate AIA activity at multiple timescales.

### Sensory pathways are integrated with metabolic state to generate local search

Previous work in insects has described the role of internal state in modulating local search behavior (Bell, 1985; Murata et al., 2017). The circuits defined here highlight the role of external sensory information in generating local search, but do not address the role of internal satiety states. To understand how internal states affect local search behavior, we inhibited feeding while animals were on food by expressing the red-shifted channelrhodopsin variant ReaChR (Lin et al., 2013) in pharyngeal muscle and using light to depolarize and paralyze the pharynx. Compared to control animals, these animals should have similar sensory experiences upon food removal but different satiety levels. We then removed these animals from food to assess local and global search behavior.

Inhibiting feeding for 10 min or 20 min prior to removal from food had little effect on the foraging sequence (Fig. 6A-B). This result suggests that the transition from local to global search that normally happens within 20 minutes is not driven by a direct input from the pharynx or by the time since feeding. On the other hand, inhibiting feeding for 45 min on food greatly reduced subsequent local search (Fig. 6C). Similarly, wildtype animals did not perform local search behavior after 45 minutes on inedible aztreonam-treated bacteria (Gruninger et al., 2008) (Fig. 6D), suggesting that the satiety state of the animal, on a longer time scale, can gate local search behavior.

**Figure 6.**
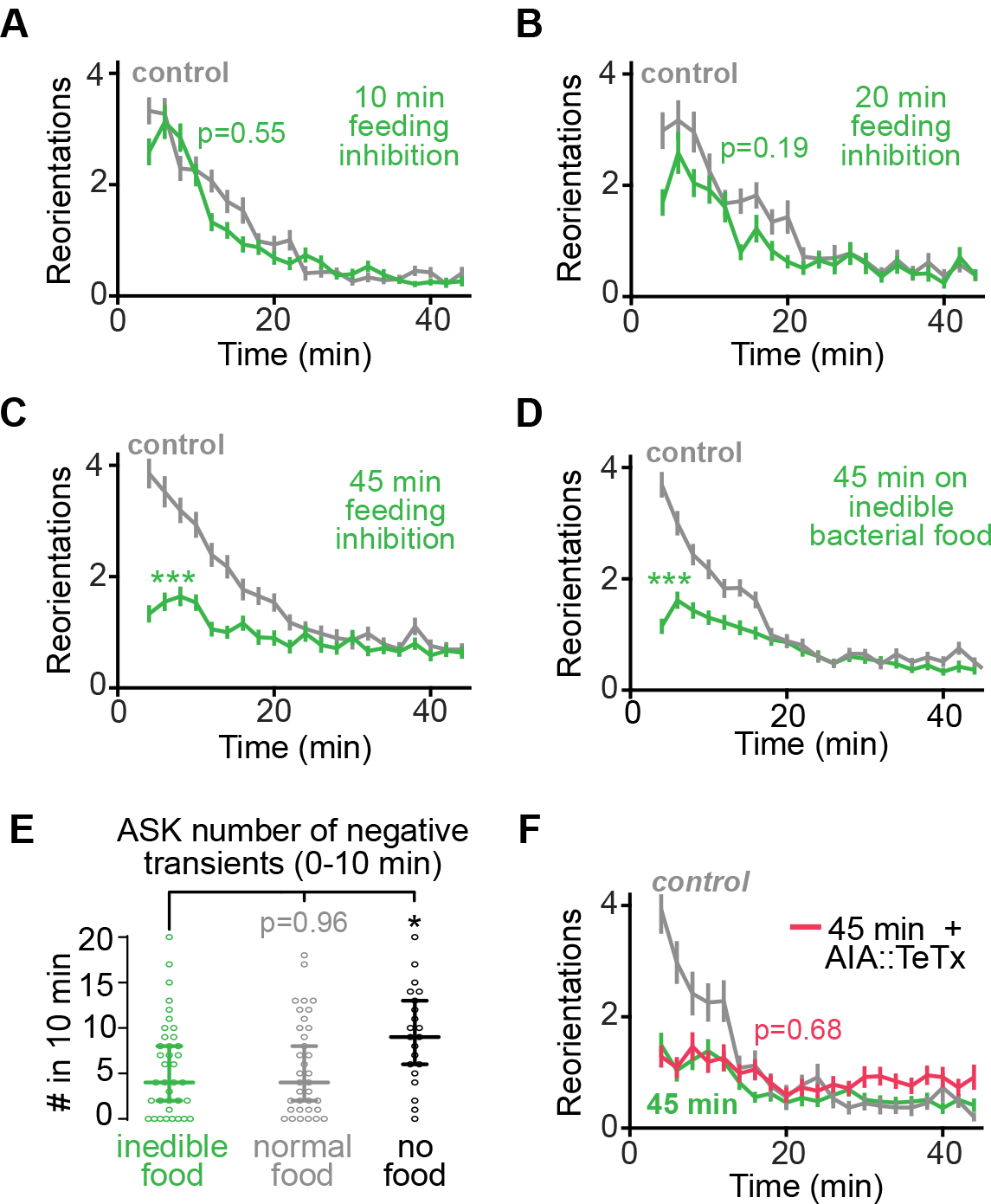
Metabolic state is integrated with sensory signals to generate local search. (A-C) Off-food foraging after feeding inhibition for various times prior to food removal. Pharyngeal pumping was inhibited by delivering light to animals expressing the red-shifted channelrhodopsin, ReaChR, in the pharynx. Control animals express ReaChR and were treated with light, but not retinal. (A) Off-food foraging after 10 min of feeding inhibition prior to food removal (control, n=37; 10 min, n=61). (B) Off-food foraging after 20 min of feeding inhibition prior to food removal (control, n=36; 20 min, n=31). (C) Off-food foraging after 45 min of feeding inhibition prior to food removal (control, n=75; 45 min, n=78). (D) Off-food foraging after 45 min on inedible aztreonam-treated bacteria. Control were animals on regular bacteria. (control, n=98; inedible bacteria, n=96). (E) ASK calcium imaging immediately (0-10min) after spending 45 min on a plate containing inedible food (n=37), normal food (n=35), or no food (n=21). Number of ASK negative transients per animal. Data presented as median with 95% confidence intervals. (F) Off-food foraging after 45 min of feeding inhibition prior to food removal + AIA silencing with tetanus toxin (TeTx) (45 min, n=47; 45 min + AIA::TeTx, n=45). All data except (E) presented as mean ± s.e.m. All p-values calculated using Wilcoxon rank sum test with Bonferroni correction. * p<0.05 ***p < 0.001.

To ask if the satiety state is encoded in the circuits defined here, we first imaged the spontaneous activity of ASK for 0-10 min after removing animals from a plate of regular food, inedible food, or no food. Animals that had spent 45 min on inedible food showed a similar number of ASK negative transients as animals that had been on regular food, whereas animals that had spent 45 min on plates without food showed a higher number of transients (Fig. 6E). These results suggest that ASK activity reports the time since the last encounter with sensory features of food, regardless of the internal satiety state.

We then asked if the satiety state interacts with MGL-1 in AIA to generate local search (Fig. 3). We inhibited feeding for 45 min in animals expressing tetanus toxin in AIA. In contrast to its strong rescue of *mgl-1* mutants (Fig. 3D) and glutamate knockouts (Fig. 4H), silencing AIA with tetanus toxin did not rescue the foraging defect resulting from feeding inhibition (Fig. 6F), indicating that the satiety state is not encoded in the activity of AIA. Together, these results suggest that satiety signals interact with parallel or downstream circuit elements to gate the ability of sensory circuits to generate local search.

## DISCUSSION

Foraging patterns incorporating local search have been described in numerous animal species (Bell, 1985; Benedix, 1993; Dias et al., 2009; Eifler et al., 2012; Gray et al., 2005; Hills et al., 2013; Lihoreau et al., 2016; Nakamuta, 1985; Papastamatiou et al., 2012; Wakabayashi et al., 2004; Weimerskirch et al., 2007). In each case, the animal performs a time-limited exploration of the region where resources were last encountered, using a set of motor programs – reversals, turns, and changes in locomotion speed – that are undirected, apparently internally generated, and relatively nonspecific, as they appear in many other spontaneous and evoked behaviors. Here, we define parallel chemosensory and mechanosensory circuit modules that each have the redundant ability to generate local search, and a third gating signal generated by an internal feeding state.

Together with previous results, our results suggest the following model for sustained local search (Calhoun et al., 2015; Gray et al., 2005; Hills et al., 2004; Wakabayashi et al., 2004). Removal from food leads to the activation of multiple sensory neurons, which release glutamate onto targets including AIA and ADE, and inhibit them through MGL-1, a G protein-coupled glutamate receptor. MGL-1 decreases the gain of AIA and ADE signaling and neurotransmitter release for a 10 to 20-minute period corresponding to local search. The reduction of either AIA or ADE activity is sufficient to modify the balance between forward and reversal states to favor reversals. AIA and ADE synapse onto the reversal-promoting AIB and AVA neurons, respectively (Fig. 7), consistent with the known antagonism between AIA (forward) and AIB (reversal) neurons and the more general antagonism between AVB (forward) and AVA (reversal) circuits (Roberts et al., 2016). After 10-20 minutes away from food, signaling from sensory neurons including ASK decreases, and the forward state is stabilized at the expense of the reversal state. After 45 minutes, a satiety signal from food is depleted or a food deprivation signal is generated to suppress local search regardless of sensory information.

**Figure 7.**
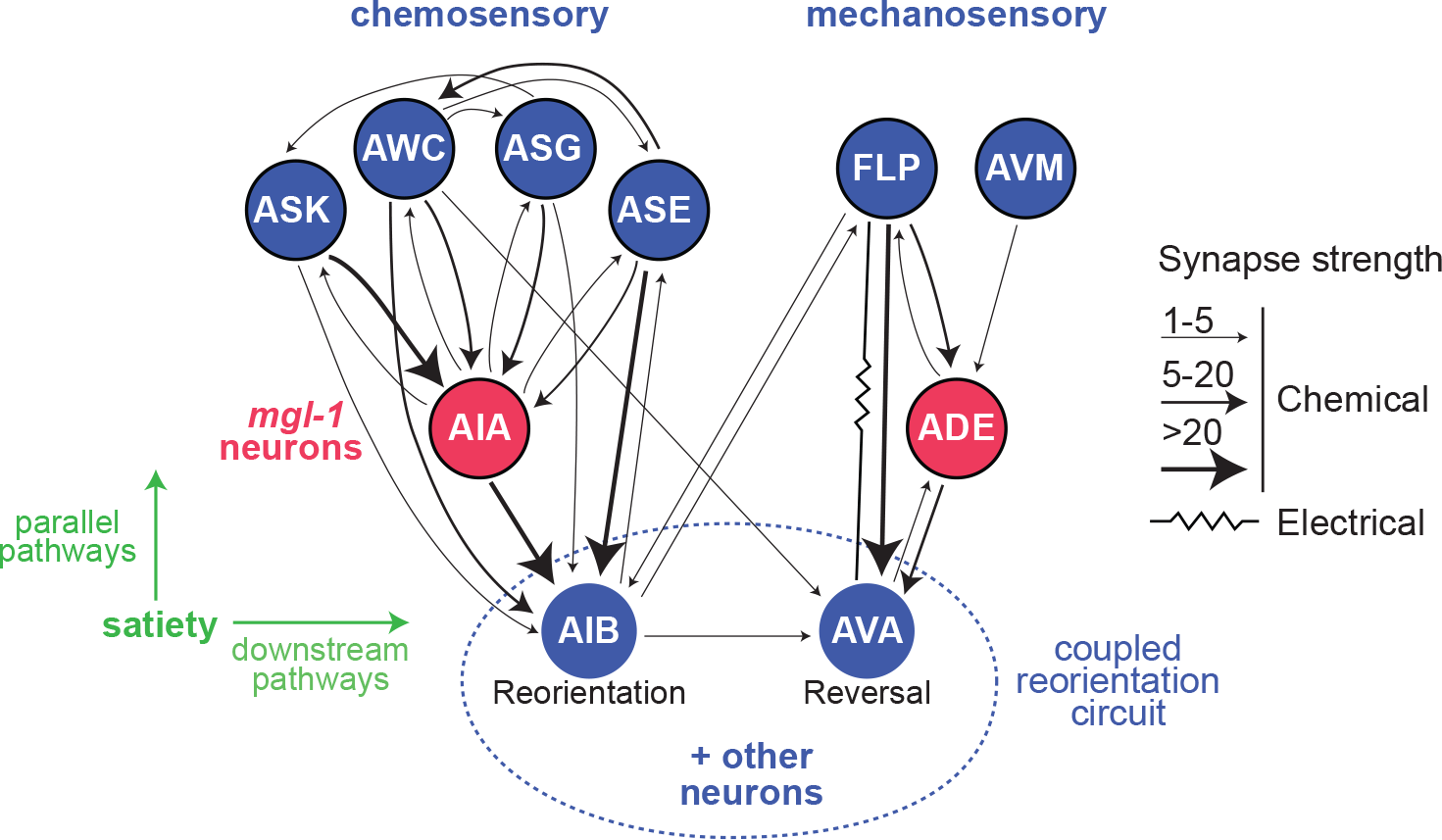
Local search circuit. Synaptic map of parallel glutamatergic modules for local search. Neurons whose activity promotes reversals are shown in blue; neurons whose activity inhibits reversals are shown in red. Arrows are weighted based on the number of chemical synapses from www.wormweb.org. The pathways consist of multimodal glutamatergic sensory neurons (blue) that converge on downstream *mgl-1-*expressing integrating neurons (red). There are no direct connections between the parallel pathways until the convergence on AIB and AVA neurons (blue), which have coupled activity that drives reversals and reorientations (coupled reorientation circuit). The satiety state of the animal can gate the behavior parallel or downstream of the circuit modules.

### Parallel sensory circuits generate multimodal control of local search

The local search circuit defined here is composed of two parallel modules: a set of glutamatergic chemosensory neurons that synapse onto the AIA interneurons, and a set of glutamatergic mechanosensory neurons that synapse onto the ADE sensory-inter neurons (Fig. 7). The chemosensory neurons detect food-related volatile odors (AWC) and amino acids (ASK), whereas the mechanosensory neurons detect touch and textures associated with food (Albeg et al., 2011; Sawin et al., 2000; Yemini et al., 2013). Each set of glutamatergic sensory neurons is predicted to inhibit AIA or ADE through the MGL-1 metabotropic glutamate receptor when animals are removed from food. Inhibition of AIA or ADE encodes a sensory food memory and allows the generation of local search. This circuit has several interesting features that shed light on multimodal sensory processing.

First, the chemosensory and mechanosensory modules that control local search are separate and parallel (Fig. 7). There are no direct connections between the identified chemosensory and mechanosensory neurons, no shared connections onto the AIA or ADE integrating neurons, and no connections between AIA and ADE in the synaptic wiring diagram of *C. elegans* (White et al., 1986). AIA and ADE primarily synapse onto AIB and AVA interneurons, respectively. AIB and AVA are two members of a coupled neuronal group that are simultaneously activated during reversal behaviors (Fig. 7) (Gordus et al., 2015; Kato et al., 2015). The coupled circuit that links AIB and AVA is the first site of convergence between the chemosensory and mechanosensory modules. Modeling suggests that stochastic fluctuations between this reversal circuit and a mutually inhibitory forward circuit may be sufficient for occasional reversals in the absence of overt sensory cues (Roberts et al., 2016); our results suggest that local search represents a change in the balance between these antagonistic assemblies.

Second, the AIA and ADE modules are each sufficient to generate full local search behavior: they are not additive. Inhibiting the function of either AIA or ADE with MGL-1 or tetanus toxin permits local search (Fig. 3). Similarly, glutamate release from either the chemosensory or the mechanosensory neurons can generate the full behavior (Fig 4). This redundant circuit organization may allow a robust innate neuronal representation of food removal through multiple sensory modalities. Indeed, there may be other sensory modules that can have the same effect. Food and satiety cues are both diverse and critical to survival, features that would favor parallel processing and redundancy in their detection. In mammals, satiety is redundantly represented by mechanosensory input from gut distension (Baird et al., 2001; Sabbatini et al., 2004) and sensory inputs associated with feeding (Chen et al., 2015). Within the mammalian brain, hunger-generating AGRP neurons also act in parallel and redundant circuits: groups of non-connected AGRP neurons project separately to distinct forebrain regions and different projections have redundant capability to elicit feeding (Betley et al., 2013). Thus, a redundant organization may be a common feature of circuits involved in survival behaviors.

### Glutamate and MGL-1 act at multiple time scales after food removal

Chemosensory neurons in the local search circuit regulate acute reorientations during behaviors like chemotaxis by activating ionotropic glutamate receptors on downstream interneurons (Hills et al., 2004; Chalasani et al., 2007). Here, we found that the same sensory neurons regulate sustained local search behavior by signaling through an inhibitory metabotropic glutamate receptor, MGL-1. Our results suggest that glutamate acts through an unknown ionotropic glutamate receptor to synchronize fast activity between ASK and AIA, and simultaneously acts over longer timescales through *mgl-1* to suppress the amplitude of AIA calcium transients (Fig. 5). Thus, the circuits that generate the fast motor components of local search may also generate the sustained search state by using different receptors.

The net effect of *mgl-1* is to inhibit its target neurons, AIA and ADE, but animals with and without *mgl-1* generate AIA calcium transients, suggesting that *mgl-1* does not eliminate neuronal excitability. *mgl-1* encodes a Group II metabotropic receptor (Dillon et al., 2006), whose mammalian homologs inhibit neurotransmitter release (Dong and Ennis, 2017; Hayashi et al., 1993); therefore *mgl-1* may decrease the synaptic output of AIA and ADE. In agreement with this possibility, inhibition of AIA or ADE synaptic release with tetanus toxin can substitute for *mgl-1* action in local search behavior (Fig. 3D-E). The nature of the relevant AIA and ADE transmitters is unknown. AIA releases acetylcholine and numerous neuropeptides, and ADE releases dopamine and several peptides including the locomotion-stimulatory peptide PDF-1. A previous area-restricted search assay identified a role for dopamine (Hills et al., 2004) that we did not recapitulate here with *cat-2* mutants, but we did identify the dopamine receptor *dop-2* in our mutant screen, suggesting that dopamine may contribute to the behavior along with other transmitters.

In addition to generating local search, *mgl-1* coordinates broader behavioral and physiological responses to food depletion. Upon food removal, *mgl-1* decreases pharyngeal pumping (feeding) (Dillon et al., 2015), promotes mobilization of fat stores downstream of TGF-β signaling (Greer et al., 2008), and modifies survival after starvation by regulating autophagy through action in AIY interneurons (Kang and Avery, 2009b). Its expression is regulated by AMPK and is dependent on satiety levels (Ahmadi and Roy, 2016). Identifying the cell type-specific readouts of MGL-1 action will provide a better understanding of conserved coordinated behavioral and physiological responses to changes in food availability.

### What dictates the duration of local search?

Theoretical models have demonstrated that local search is the most successful strategy when animals have information about food availability, such as a recent food encounter, and that a global search is most successful when animals need to locate random food sources with no prior information (Calhoun et al., 2014; Salvador et al., 2014). The circuits described here can encode the recent food encounter through sensory cues, and thereby drive an active local search behavior that decays over time to a global search behavior. One open question that remains is the nature of the internal clock that sets the duration of local search. Our results suggest that internal clocks use a suite of physiological and molecular mechanisms to represent food over time.

The timing of AIA and ADE inhibition by glutamate and MGL-1 represents sensory history, but it can be separated from the internal clock for local search duration. Tetanus toxin inhibits AIA and ADE synaptic release across life but does not affect the transition from local to global search (Fig. 3D-E). This result strongly argues that structured AIA and ADE activity alone do not define the duration of local search.

Nutritional cues act over slower timescales. Inhibiting feeding for 10 min or 20 min before food removal did not affect the timing of local search, which is largely complete in 20 min after food removal (Fig. 6A-B). Nonetheless, sustained inhibition of feeding for 45 minutes eliminated all local search behavior (Fig. 6C-D). Unlike *mgl-1* or glutamate knockout, the effect was not rescued by introducing tetanus toxin into AIA (Fig. 6F). The all-or-none and delayed nature of this satiety effect suggests that metabolic cues from feeding regulate local search downstream or in parallel to the sensory cues represented in the *mgl-1* circuit (Fig 7). In this formulation, internal satiety signals gate the ability of sensory information to generate behavior. The role of satiety in sensory food information may allow animals to represent food not only by its multiple sensory modalities, but also by its nutritional quality.

Together with these mechanisms, neuropeptides released from sensory neurons are attractive candidates to set the duration of local search. Local search is affected after laser ablation of either AWC or ASK sensory neurons (Gray et al., 2005), but our results show that glutamate release from either neuron can be eliminated with less effect (Fig. 4C-D). Cotransmitters such as neuropeptides could explain the difference between glutamate release and neuronal ablation, although other explanations are also possible. Most of the sensory neurons in the local search circuit release FMRF neuropeptides, which are related to mammalian NPY peptides (Li and Kim, 2008), and several neuropeptides and G protein-coupled neuropeptide receptors affect local search (Campbell et al., 2016; Chalasani et al., 2010). In the rodent hypothalamus, NPY cotransmitter release converts transient GABA-dependent synaptic effects into sustained feeding behavior (Chen et al., 2016). By analogy, *C. elegans* sensory neuropeptide release could act with glutamate to generate sustained local search behaviors.

The temporal heterogeneity of fast glutamate receptors, slower MGL-1 and neuropeptide signaling, and nutritional input may store food memory in multiple clocks. Ultimately, this memory must be manifested in neuronal activity. One representation of food history that merits further study is the spontaneous activity of the ASK neuron, which changes over a time frame that matches the transition from local to global search (Fig. 5). A recent study of *C. elegans* brain states at long times after food removal reported changes in the endogenous activity in ASK, in agreement with our results (Skora et al., 2018). Identifying the sources of slowly-changing spontaneous activity in ASK may provide further understanding of how behavioral states are sustained and terminated.

## METHODS

### Nematode Culture

Animals were grown at room temperature (21-22°C) on nematode growth media (NGM) plates seeded with *E. coli* OP50 bacteria (Brenner, 1974). Wildtype animals were *C. elegans* Bristol strain N2. All mutant strains tested were backcrossed to wildtype animals to reduce unannotated background mutations. Mutants strains used for the candidate genetic screen are listed in Table S1. CRISPR-generated strains and transgenic strains are listed in Table S2. Mutant strains used in candidate screen were generated by other groups using random mutagenesis and selected by us based on the annotated mutations. Transgenics were generated by microinjection of a transgene derived from wildtype strain N2, a fluorescent co-injection marker (*myo-2::mCherry, myo-3:mCherry, elt-2::nls-GFP, elt-2::mCherry*), and empty pSM vector to reach a final DNA concentration of 100 ng/μL.

Transgenic and mutant strains were always compared to matched controls tested in parallel. Each experiment included 12-18 animals. For the candidate genetic screen, each mutant strain was tested in six different experiments done on two different days. All other strains were tested in 4-8 experiments done on at least two different days.

### CRISPR/Cas9-generated mutant strains

CX1030 *npr-9(ky1030)* X. *ky1030* is a CRISPR/Cas9 induced single nucleotide deletion in the second exon of *npr-9*. The resulting sequence is TGGTAATGCT<u>C</u>TGGTGGTGAT (deleted nucleotide is underlined). To generate the strain, we used a co-CRISPR protocol (Arribere et al., 2014). Young hermaphrodites were injected with a mix of plasmids encoding Cas9, a gRNA targeting *rol-6*, a gRNA targeting *npr-9*, and a ssDNA repair template that induces a dominant *rol-6(su1006)* mutation. F1 animals were isolated to individual plates based on their roller phenotype, allowed to lay eggs, and then screened for a target mutation by Sanger sequencing. F2 animals were isolated to find homozygotes for the target mutation. After inducing the target mutation, animals were backcrossed to wildtype twice.

CX1037 *mgl-1(ky1037)* X. *ky1037* is a CRISPR/Cas9 induced indel that results in a frameshift before the first transmembrane domain of *mgl-1*. Deletion is (G), insertion is (TTGTGTGGTTGTGTGGTTGTGGTTGTGTGTTGT). Resulting sequence is TGGAGCAACG<u>TTGTGTGGTTGTGTGGTTGTGGTTGTGTGTTGT</u>TGGTGGTTCT (insertion is underlined). We used the co-CRISPR protocol as described above (Arribere et al., 2014). pJA42 was a gift from Andrew Fire (Addgene plasmid # 59930)

### CRISPR/Cas9 editing of the endogenous *eat-4* (VGLUT1) locus

To generate a strain that would allow us to knock out endogenous glutamate release in a cell-specific manner, we performed two successive edits on the endogenous *eat-4* locus.

First, we used the co-CRISPR method described above (Arribere et al., 2014) to insert an FRT sequence (GAAGTTCCTATTCTCTAGAAAGTATAGGAACTTC) immediately before the start codon of *eat-4*. We injected the mix described above with an additional ssDNA repair template consisting of the FRT sequence with 35 bp homology arms on each side. The resulting edited genomic sequence is:

*TCATCATCATTTTCAGAAACC*<u>GAAGTTCCTATTCTCTAGAAAGTATAGGAACTTC</u>***ATG**TCGTCATGGAACGAaGC* (FRT insertion is underlined; *eat-4* locus is in italics; bolded ATG is the start codon of *eat-4*; lowercase *‘a’* is a silent mutation induced to remove PAM site after successful homologous recombination).

Second, we used the CRISPR/Cas9 SapTrap method (Schwartz and Jorgensen, 2016) to insert *let-858 3̍-UTR::FRT::mCherry* immediately after the endogenous *eat-4* stop codon. SapTrap is a modular plasmid assembly approach that produces a single plasmid vector containing both a gRNA transcript and a repair template. We mutated some of the modular components used to assemble the repair template as follows: for pMLS279 (*FRT-let-858 3̍-UTR-FRT*) we deleted the 5̍ FRT site (new plasmid is pALC01), and for pMLS291 (mCherry with syntron embedded inverted floxed *Cbr-unc-119*) we added a stop codon at the end of the mCherry coding region (new plasmid is pALC02). We cloned these two edited plasmids along with 60 bp homology arms and the gRNA into the pMLS256 destination vector, producing a combined plasmid vector (pALC03). Young *unc-119(ed3)* hermaphrodites were injected with the combined plasmid vector, a Cas-9 expression vector, and fluorescent co-injection markers. F1 animals were picked 10 days after injection based on rescue of the *unc-119* phenotype, and the insertion was confirmed by PCR and Sanger sequencing. These edited animals were then injected with *peft-3::nCre* (pDD104) to excise the syntron embedded, inverted floxed *Cbr-unc-119*. F1 animals were selected based on the reappearance of the *unc-119* phenotype. This line was backcrossed once to wildtype animals to remove the *unc-119(ed3)* mutation, and an additional three times to remove any off-target CRISPR mutations. The resulting strain is CX17461.

Nuclear localized flippase (nFLP) was subcloned from pMLS262 (snt-1::2xNLS-FLP-D5) into pSM using Gibson assembly (New England Biolabs). A plasmid containing nFLP under cell specific promoters was injected into the *eat-4* edited strain (CX17461) by standard gonadal microinjection. Successful glutamate knockout in the target cells was confirmed by mCherry expression which was visualized using a Zeiss Axio Imager.Z1 Apotome microscope with a 40x objective.

For all glutamate knockout experiments (Fig. 4), ‘control’ animals are the *eat-4* edited strain, and *‘mgl-1’* are *mgl-1(ky1037) eat-4* edited strain.

pMLS279 (Addgene plasmid # 73729), pMLS291 (Addgene plasmid # 73724), pMLS256 (Addgene plasmid # 73715), and pMLS262 (Addgene plasmid # 73718) were gifts from Erik Jorgensen.

### Identification of *mgl-1(ky1060)* allele

We prepared genomic DNA from strain CX17083 using standard phenol/chloroform extraction protocols. Sequencing was conducted at the Rockefeller High-Throughput Sequencing Facility, where a DNA library was prepared using TruSeq Nano DNA kit (Illumina) and sequenced in an Illumina HiSeq 2500 System.

Deletions were identified using modifications to a previously described approach (McGrath et al., 2011). Candidate deletions were first identified using ‘chimeric’ reads as identified by bwa (i.e. reads that included the SA tag) where each alignment mapped to the same chromosome and strand (Li and Durbin, 2010). These reads were used to infer the breakpoints and insertion sequence of a candidate deletion. Candidate deletions that were also present in N2 sequenced controls were excluded as likely errors in the reference sequence. Each candidate deletion was then genotyped by collecting all the reads with primary alignments that fell within 10 bp of the candidate deletion and a mapping quality score greater than 10. These reads were realigned to both the reference and the candidate deletion sequence using a striped Smith Waterman Alignment implemented in the scikit-bio Python library (http://scikit-bio.org/). This analysis identified a homozygous 225 bp deletion in the *mgl-1* gene that was verified using Sanger sequencing.

The identified *mgl-*1 deletion is:

ATCTGCTCAACGACCAAGATTCATATCTCCCATCTCTCAGgtgagctccggtgacaagccaa cggaagtacactatttatagGTTGTCATGACTGCAATGCTAGCCGG AGTACAATTGATCGGAAGTCTTATTTGGCTGTCAGTAGTGCCACC AGgtaaattggctatttatgaagtgatgtctgagtaatttttag GTTGGAGACACCACTACCCCACCAGGGACCAGGTGGTTTTAACTTGTAATGT TCCTGACCATCACTTTTTGTATTC (deletion underlined, exons in uppercase, introns in lowercase)

### Intersectional Cre/Lox *mgl-1* rescue

To rescue *mgl-1* in subsets of its endogenous expression pattern, we employed an intersectional transgene strategy depicted in Figure 3A. The inactive *mgl-1* genomic fragment plasmid was constructed using pSM-inv[sl2-GFP] as the backbone (Flavell et al., 2013). A portion of a *mgl-1* rescue genomic fragment sequence was cloned into pSM-inv[sl2-GFP] in its correct orientation (obtained using forward primer: 5′- GTAAGGTATGTTTTTATTTTCCAAC -3′ and reverse primer: 5′- TAGAACAGACAAACATATTTGAC -3′, and cloned in before the first Lox2272 and LoxP sites). The remaining *mgl-1* fragment sequence was cloned into pSM-inv[sl2-GFP] in an inverted orientation (obtained using forward primer: 5′-TCATAAGAAAGTATCGTGAGC -3′ and reverse primer: 5′- CATTATGGCGTATGATGGG -3′, cloned in after the inverted sl2-GFP and before the inverted LoxP and Lox2272 sites). The cloning was done via Gibson assembly (New England Biolabs).

To rescue *mgl-1* in subsets of its endogenous expression pattern, we first injected the partially inverted plasmid described above into *mgl-1(ky1037)* animals using standard microinjection protocols. The resulting line is labelled ‘no Cre’ in Fig. 3B-C and Fig. S4A-B. Subsequently, we injected plasmids expressing nls-Cre under cell specific promoters into this ‘noCre’ line. This led to Cre-mediated inversion and reconstitution of the *mgl-1* rescue fragment in subsets of cells, which was confirmed by GFP expression visualized using a Zeiss Axio Imager.Z1 Apotome microscope with a 40x objective

### Foraging assay and quantification

Bacterial food lawns were made by seeding NGM plates with a thin uniform OP50 lawn (OD~0.5) 16 hours before the assay. The lawn covered the entire plate to eliminate effects of animals exploring the lawn edge (Calhoun et al., 2015), and contained a filter paper barrier soaked in 20 mM CuCl_2_ that prevented the animals from leaving a 5 cm × 5 cm region on food. On the assay day, 10-15 adult hermaphrodites were first transferred for 45 min to this standard food lawn. To study off-food foraging, animals were transferred from the standard food plate to an unseeded NGM plate, allowed to crawl at least five body lengths to clean off excess food, and transferred to the assay plate which consisted of a large NGM plate with a circular filter paper barrier (~80 cm^2^) soaked in 20 mM CuCl_2_ to restrict animals to the recorded area. Their behavior was recorded for 45 min, starting four minutes after the initial food removal, using a 15 MP PL-D7715 CMOS video camera (Pixelink). Frames were acquired at 3 fps using Streampix software (Norpix). Individual worm trajectories were analyzed using custom Matlab (Mathworks) software, as previously described (Pokala et al., 2014). We were able to track the behavior of some individuals for the entire 45 min of food. When collisions occurred, however, the data points around the collision were excluded (to ignore reorientations associated with collisions) and sometimes we were unable to link tracks after the collision, resulting in separate tracks before and after collisions. These track fragments were included in average quantifications.

To quantify reorientations frequencies, we counted the number of reorientations that each animal performed in 2-minute time windows and divided it by the number of animals tracked in that time window. We only counted animals that we were able to track in the entire 2-min time window. The plots show the mean number of reorientations in 2-min time windows. In all figures (except Fig. S1), we quantified the number of time-dependent reorientations only (reorientations shown in Fig. S1A-C).

To calculate Mean Square Displacement (MSD) (Fig. 1B), we used the centroid x-y position of each animal. MSD was defined as the square of the distance travelled in one minute. For each animal, we first we calculated the MSD for each frame by measuring the squared of the distance travelled from 30 sec before the frame to 30 sec after the frame. We then calculated the average MSD in two-minute time windows for each animal (the average MSD in 360 frames = 2 min).

For the food concentration experiments (Fig. 1D), the usual OP50 culture (OD~0.5) was diluted 10x or 100x in LB before seeding on NGM plates and growing for 16 hours overnight. The off-food experiment was the same as described above.

### Histamine silencing experiments (Pokala et al., 2014)

Neurons expressing histamine-gated chloride channels (HisCl1) were silenced by adding histamine to NGM assay plates. 1 M histamine-dihydrochloride (Sigma-Aldrich) in deionized water was filtered and stored at −20°C. NGM histamine plates were made by adding 1M histamine to NGM solution (50-55°C) to a final concentration of 50 mM histamine. Plates were stored at 4°C and used within a week. Controls included: (1) animals expressing the HisCl1 tested on normal plates and (2) animals with no transgene tested on histamine NGM plates.

### Calcium imaging

Transgenic animals expressing GCaMP5A (Akerboom et al., 2012) in ASK (*sra-9::GCaMP5A*), in AIA (*gcy-28d::GCaMP5A*), or in both neurons (crossing the two single neuron GCaMP5A lines), were generated in a wildtype background and crossed into *mgl-1 or eat-4* if indicated. Adult animals were first transferred to a standard food lawn for 45 min prior to imaging. Animals were then removed from the lawn, washed in NGM buffer to remove any food, and loaded into a custom PDMS microfluidic chamber in NGM buffer, where they were restrained but not paralyzed (Chronis et al., 2007). Animals received a constant flow of NGM buffer through the experiment. Imaging began 6 min after removal from food (time=0 min). Animals were imaged for 10 min initially (0-10 min), left in the device for 30 min with the light off, and imaged again for another 10 min (40-50 min). Calcium fluorescent signals were acquired at 5 frames per second through a 40x objective on an upright Axioskop 2 microscope (Zeiss) with an iXon3 DU-897 EMCCD camera (Andor), using Metamorph Software (Molecular Devices). Custom Image J and MATLAB software were used to track, quantify, and analyze neuronal activity. ASK imaging data was bleach-corrected. AIA did not show significant bleaching.

For ASK, the maximum baseline fluorescence in 1-minute intervals, F_max_ (95th percentile value), was used to calculate the change in fluorescence for each frame in the interval (ΔF = F-F_max_), and subsequently normalized to F_max_ (ΔF/F_max_). Changes in fluorescence observed were spontaneous decreases from the ASK baseline fluorescence (negative transients). To characterize negative transients, we wrote custom MATLAB software where we defined parameters to identify transients (start/end of transients = −0.1ΔF/F_max_, minimum transient amplitude = −0.3ΔF/F_max_, minimum transient duration = 3 sec). Additionally, we used the MATLAB function *findpeaks* to identify complex transients which consist of multiple local minima within one transient (arrowheads in Fig. 5A). Parameters were optimized to match manually assigned negative transients. For transient number and amplitude quantifications we measured all local minima within a complex transient. For transient duration quantifications we calculated the duration of the entire complex transient. For *eat-4* analysis, we used the same parameters used in wildtype data.

For AIA, the minimum baseline fluorescence, F0, was calculated using the MATLAB function *statelevels* which divides the data into two histograms and computes the mode of the lower histogram. F_0_ was used to calculate the change in fluorescence for each frame (ΔF = F− F_0_), and subsequently normalized to F_0_ (ΔF/ F_0_). AIA shows a low baseline fluorescence with spontaneous positive transients. To characterize positive transients, we used the same procedure described above (parameters: start/end of transients = 0.1ΔF/F_0_, minimum transient amplitude = 0.3ΔF/F_0_, minimum transient duration = 2 sec). Parameters were optimized to match manually assigned transients. For transient number and amplitude quantifications we measured all local maxima within a complex transient. For transient duration quantifications we calculated the duration of the entire complex transient. For *mgl-1* and *eat-4* analysis, we used parameters used in wildtype data.

To align ASK traces to AIA positive transient onset in the simultaneous imaging experiments (Fig. 5G), we first identified the start time of each AIA transient, and then extracted ASK frames from 10 sec (50 frames) before the start time until 30 s after the start time. We then averaged all the extracted ASK frames.

To calculate correlations, we first z-scored each trace and then computed the autocorrelation, or the cross-correlation between ASK and matching lagged AIA traces, using the MATLAB function *xcorr*.

### Optogenetic neuronal manipulations

To silence neurons optogenetically we expressed the light-gated anion channel, *Guillardia theta* anion channel rhodopsin 2 (*Gt*ACR2). A codon-optimized *Gt*ACR2 was synthesized based on the *Gt*ACR2 ORF (Govorunova et al., 2015) and cloned into the pSM vector backbone via Gibson Assembly (New England Biolabs).

To express *Gt*ACR2 selectively in glutamatergic sensory neurons that synapse onto AIA and ADE (the same neurons where glutamate knockout leads to local search defect, Fig. 4), we employed an inverted Cre-Lox recombination strategy. A floxed *Gt*ACR2 cDNA was placed in an inverted orientation under the *tax-4* (chemosensory subset) and *mec-3* promoters (mechanosensory subset), and subsequently activated in the relevant cells by Cre expression under the *eat-4* promoter (glutamate specific).

16 hours before the experiment, L4 animals expressing *Gt*ACR2 were picked to NGM plates seeded with OP50 containing 12.5 μM all-trans retinal (Sigma). Control animals, which also expressed *Gt*ACR2, were transferred to NGM plates seeded with OP50 but no retinal. On the experimental day, 15-20 animals were first transferred to a standard food lawn (with or without retinal) for 45 min, and then transferred to the large off-food assay plate (~80 cm^2^) where their behavior was recorded as described above. During video recording, green light (~45 μW/mm^2^) was delivered at 10 Hz (50% duty cycle) using a Solis High Power LED (ThorLabs) controlled with custom MATLAB software.

For the sensory silencing experiments, we delivered four 30 s light pulses with 2 min in between each pulse. The pulses began 2 min after initiating recording. To analyze the effect of the neuronal manipulation, we calculated the fraction of animals reorienting in each frame. We aligned all pulses and averaged the behavior for ten different experiments (40 light pulses).

### Pumping inhibition experiments

16 hours before the experiment, L4 animals expressing *myo-2::ReaChR* were picked to NGM plates seeded with OP50 containing 50 μM all-trans retinal (Sigma). Controls animals, which also expressed *myo-2::ReaChR*, were transferred to NGM plates seeded with OP50 but not retinal. On the experimental day, 15-20 animals were transferred to a standard food lawn (without retinal) for 45 min, and green light was delivered (10 Hz, ~45 μW/mm^2^) for different amounts of time (10, 20, 45 min) prior to food removal. Animals were recorded on food to monitor their speed. Successful feeding inhibition was confirmed by a sustained increase in locomotion speed on food (increased roaming). The off-food experiment was the same as described above.

### Inedible bacteria experiments

Inedible bacteria were generated using a modified protocol developed by Gruninger et. al. In this method, bacteria were treated with a low concentration of aztreonam which prevents septum closure during division, resulting in long chains of undivided bacteria. It has been shown that these inedible bacteria have similar sensory properties to regular bacteria, but that they cannot be ingested by *C. elegans* (Gruninger et al., 2008).

To make these inedible bacteria, OP50 cultures were grown overnight in LB at 37°C. The next day this culture was diluted 5x in LB with aztreonam (Sigma) at final concentration of 10 μg/ml, and grown overnight at 37°C. The next day, 16 hours before the assay, this culture was seeded on NGM plates with 10 μg/mL aztreonam. Control lawns were made the same way, but without adding aztreonam. Lawns were thin, and uniform as described above, and covered the entire plate. Aztreonam and control plates were made fresh for every experiment.

### Statistical Methods

Differences in local search behavior between groups was assessed by counting the number of reorientations that each animal in each group performed 2-12 min after food removal. Statistical significance was assessed using a Wilcoxon rank sum test with Bonferroni correction. The Wilcoxon signed rank test was used to compare matched data (imaging experiments, same animal early vs late). The Kolmogorov-Smirnov test was used for comparing cumulative probability distributions.

## AUTHOR CONTRIBUTIONS

A.L.C. and C.I.B. designed experiments, A.L.C. conducted experiments, N.P. developed automated behavioral tracking code, A.L.C. and A.S. generated the *eat-4* CRISPR glutamate knockout line, P.T.M. mapped the *mgl-1* deletion in CX17083, S.W.F. developed optogenetic feeding inhibition protocols. A.L.C and C.I.B. analyzed and interpreted results, and A.L.C. and C.I.B. wrote the paper.

## ACNOWLEDGMENTS

We thank the Caenorhabditis Genetics Center (NIH P40 OD010440) and the Million Mutation Project for strains; E. Scheer and P. Kidd for advice and discussions; S. Stern, S. Levy, M. Dobosiewicz, and J. Jimenez for comments on the manuscript. A.L.C. was supported by an F31 Predoctoral Fellowship from the National Institute of Mental Health of the National Institutes of Health under award number F31MH108325, and by a Medical Scientist Training Program grant from the National Institute of General Medical Sciences of the National Institutes of Health under award number T32GM007739 to the Weill Cornell/Rockefeller/Sloan Kettering Tri-Institutional MDPhD Program. This work was supported by the Howard Hughes Medical Institute, of which C.I.B. was an investigator.

## Supplemental Figure Legends

**Figure S1.**
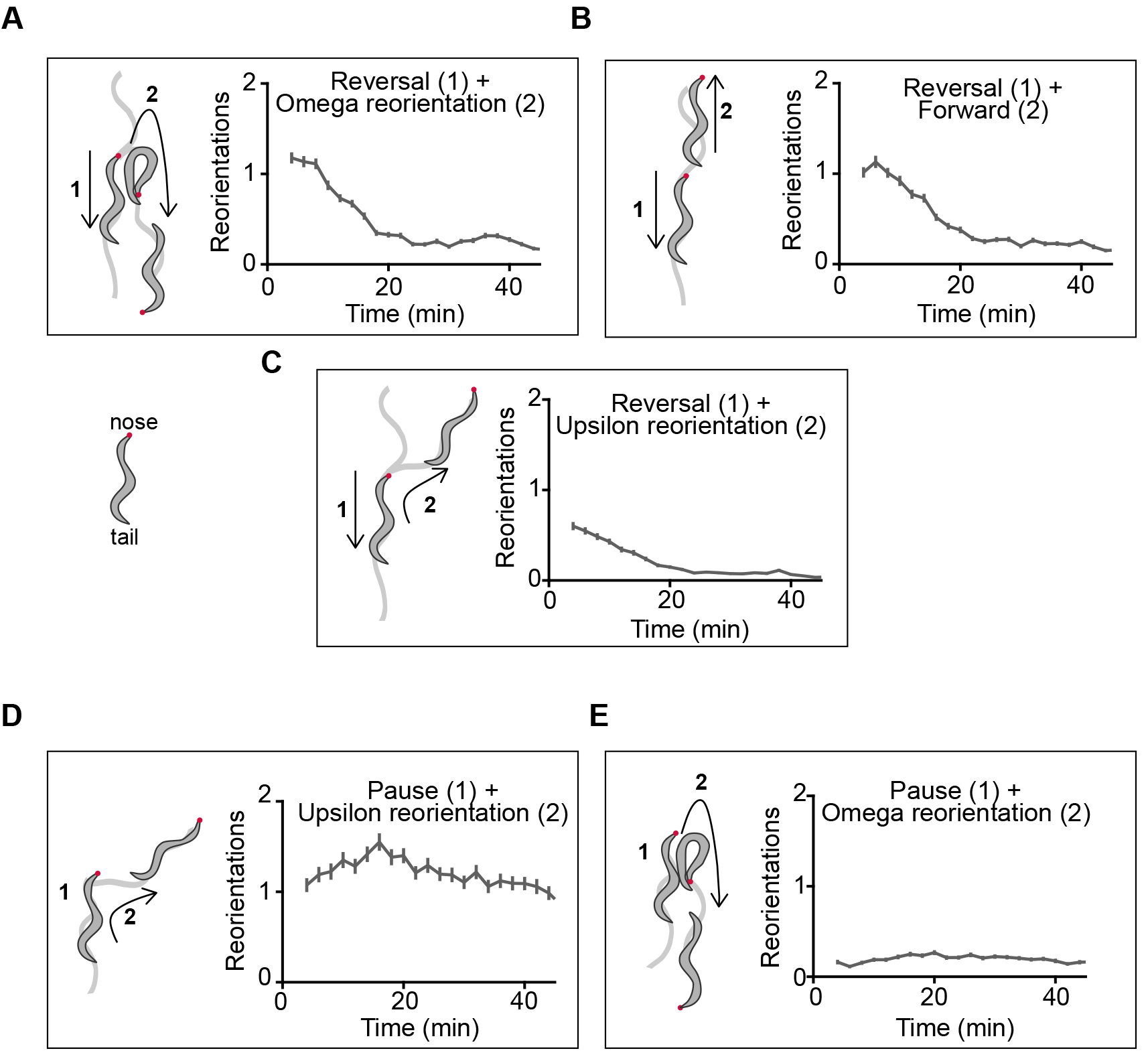
Reorientation categories fall into time-dependent or time-independent, Related to Figure 1. (A-C) Reorientations whose frequency is dependent on the time since food removal. All include an initial reversal. (A) Reversal + deep bend (Omega). (B) Reversal only. (C) Reversal + shallow bend (Upsilon). (D-E) Reorientations whose frequency is not dependent on time since food removal. (D) Pause + Upsilon. (E) Pause + Omega. Data also includes animals that show an abrupt Upsilon or Omega during forward movement, without a pause.

**Figure S2.**
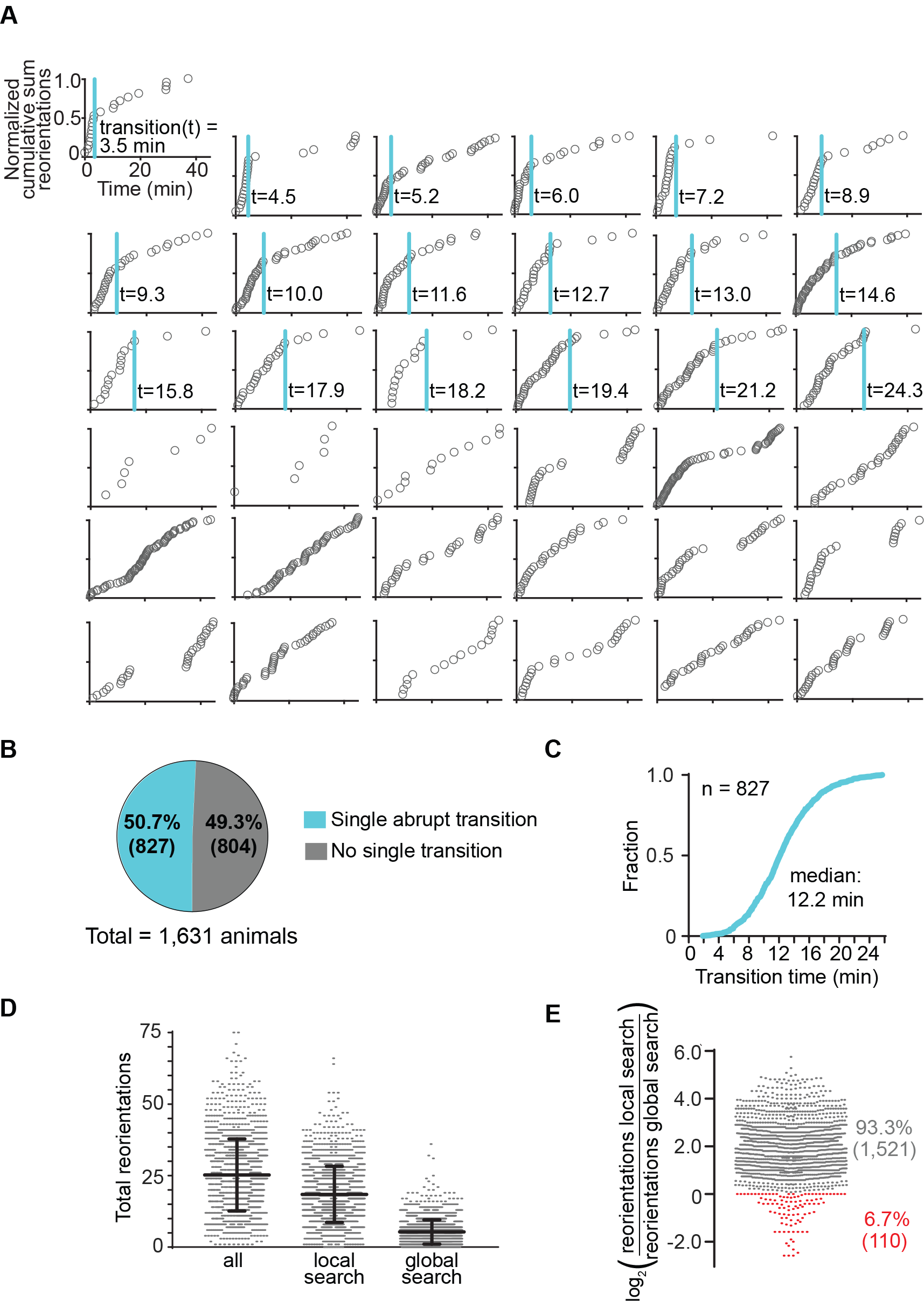
Reorientation dynamics in individual animals, Related to Figure 1. (A-C) Off-food reorientation dynamics for 1,631 individual animals from 198 different experiments collected over two years. (A) Examples of individual animals. Normalized cumulative sum of reorientations after food removal for 36 individual animals. Each circle represents a reorientation. Top 18 panels showed a single abrupt behavioral transition (marked with blue line) recognized as a change in the slope of the cumulative sum; bottom 18 panels did not. Each individual animal was analyzed by: (1) using the MATLAB function *findchangepts* to find a potential transition time. This function divides each trace into two regions that minimize the sum of the residual squared error of each region from a local linear regression, and then (2) visually inspecting the potential transition time to determine if it corresponded to a single transition. (B) Percentage of animals that performed abrupt transitions vs. non-abrupt transitions based on the method described above (n = 1,631). (C) Cumulative distribution of transition times for animals that showed a single abrupt transition (n=827). The abrupt transition time corresponds to the end of local search. Previous studies had reported that either all animals showed an abrupt behavioral transition (Calhoun et al., 2014), or that no animals showed abrupt transitions (Klein et al., 2017). Here, by looking at many animals, we show that both types of dynamics exist within the population. (D) Total number of reorientations of individual animals during the entire 45 min off food, during the first 20 min (local search), and during the last 20 min (global search). There is high variability in the reorientation rates of individual animals (n=1,631). (E) log2 (number of reorientations in first 20 minutes / number of reorientations in last 20 min) for each individual animal. 93.3% of animals perform more reorientations early than late after food removal (1,521/1,631; gray dots), suggesting that local search is a robust response to a recent food encounter. 6.7% of animals perform the same number or fewer reorientations early than late after food removal (84/1,631; red dots).

**Figure S3.**
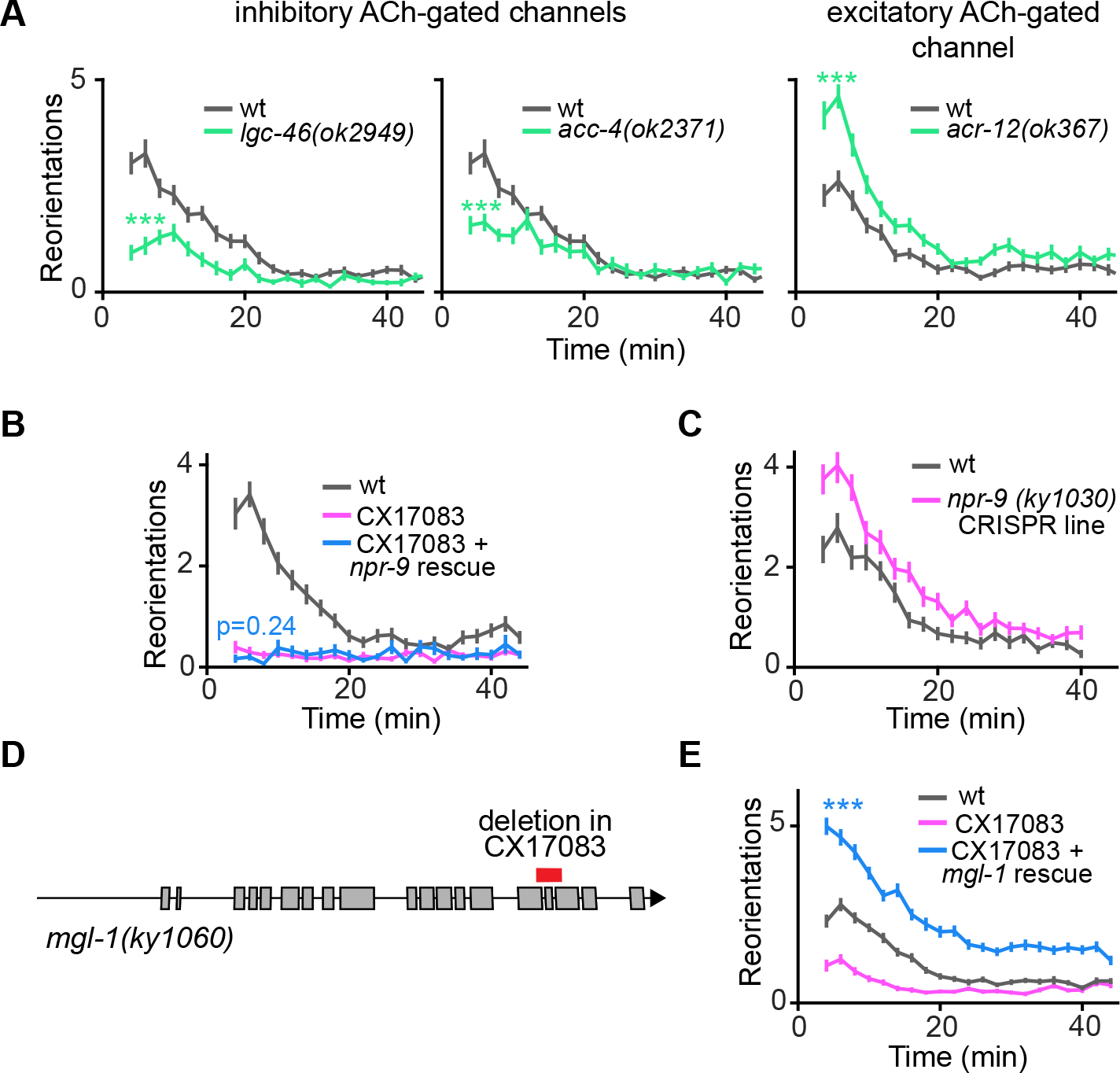
Acetylcholine receptor mutants, *npr-9* rescue experiments, and *mgl-1* mapping and rescue, Related to Figure 2. (A) Off-food foraging in mutants for the inhibitory acetylcholine receptors, *lgc-46* and *acc-4*, and for the excitatory acetylcholine receptor, *acr-12*. (wt_*lgc-46,acc-4*_, n=55; *lgc-46*, n=51; *acc-4*, n=55; wt_*acr-12,*_, n=56; *acr-12,* n=61). All three are ligand-gated ion channels that are broadly expressed in interneurons and cholinergic motor neurons. (B) Off-food foraging defect in CX17083 (pink) is not rescued by a fosmid containing an *npr-9* genomic fragment (blue) (CX17083, n=51; *npr-9* fosmid, n=29). (C) Off-food foraging of the CRISPR/Cas9 generated *npr-9(ky1030)* mutant does not recapitulate the phenotype in CX17083. (D) *mgl-1* deletion in CX17083, identified by whole genome sequencing. Deletion spans three exons and is predicted to affect transmembrane domains in all *mgl-1* isoforms. (E) Foraging defect in CX17083 (pink) is rescued by a PCR product encompassing the *mgl-1* genomic region (blue) (CX17083, n=87; *mgl-1* PCR rescue, n=88). All data presented as mean ± s.e.m. p-values calculated using Wilcoxon rank sum test with Bonferroni correction ***p < 0.001. A previous study (Campbell et al., 2016) reported that *npr-9(tm1652)* has a decreased reorientation baseline off-food compared to wildtype. The results shown in panels B-E suggest that this effect may be due to the background *mgl-1(ky1060)* mutation. The same study reported that *npr-9* overexpression suppresses reorientations, consistent with the reciprocal increase in reorientations we observed in the CRISPR-generated *npr-9(ky1030)* loss-of-function mutant.

**Figure S4.**
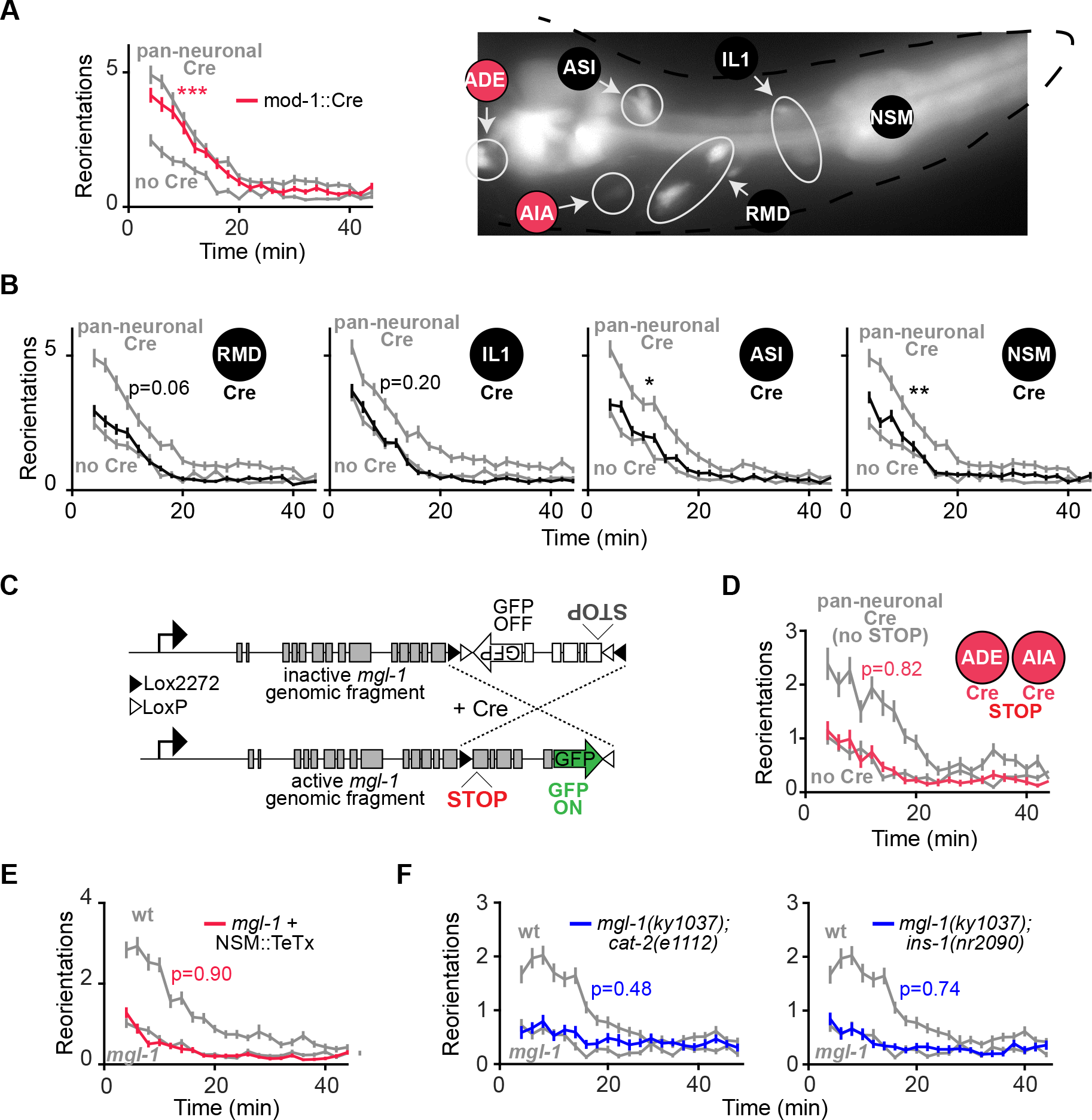
*mgl-1* cell specific rescues, transgene controls, and tetanus toxin control, Related to Figure 3. (A) *mgl-1* expression in subsets of its endogenous expression pattern using an intersectional transgene approach (Fig. 3A). Animals that only express the inverted transgene are labelled ‘no Cre’. (A) Activating the *mgl-1* inverted transgene in AIA, ADE, RMD, IL1, ASI and NSM using *mod-1::nCre* rescues local search behavior (no Cre, n= 72; *mod-1::nCre* rescue, n=80). Left: Behavioral results. Right: GFP expression in rescued animals reporting intersection of *mgl-1* and *mod-1* expression. ADE expression of *mgl-1* was not reported in previous reporter fusions with upstream regions but is seen in our genomic construct that includes introns. ADE expression of *mgl-1* is supported by single-cell transcriptomic analysis that found *mgl-1* expressed in a cluster of dopaminergic cells that includes ADE (Cao et al., 2017). (B) Minimal rescue by expression of *mgl-1* in RMD, IL1, ASI or NSM. (no CreRMD, n= 72; RMD rescue, n=76; no Cre_IL1_, n= 68; IL1 rescue, n=82; no Cre_ASI_, n= 78; ASI rescue, n=89; no Cre_NSM_, n= 72; NSM rescue, n=69). ASI and NSM rescue are statistically different from ‘no Cre’ but the magnitude of rescue is very small, so it may not be biologically meaningful. Consistent with this, silencing NSM with TeTx does not rescue *mgl-1* (below, E). (C) Control Cre-Lox *mgl-1* transgene, which when inverted has two adjacent nonsense mutations. (D) Reconstituting nonsense *mgl-1* transgene in AIA and ADE does not rescue local search. Animals that only express the inverted nonsense transgene are labelled ‘no Cre’ (no Cre, n=66; AIA+ADE nonsense rescue, n=62). (E) No rescue of *mgl-1* mutant phenotype by expression of tetanus toxin light chain (TeTx) in NSM (*mgl-1*, n=94; NSM::TeTx *mgl-1*, n=93). (F) *Left:* Off-food foraging for *mgl-1;cat-2* double mutants is indistinguishable from *mgl-1* single mutants, suggesting that *mgl-1* does not only act by decreasing dopamine release from ADE. *Right:* Off-food foraging for *mgl-1;ins-1* double mutants is indistinguishable from *mgl-1* single mutants, suggesting that *mgl-1* does not only act by decreasing *ins-1* release from AIA. (*mgl-1,* n=78; *mgl-1;cat-2*, n=80; *mgl-1;ins-1,* n=79). All data presented as mean ± s.e.m. p-values calculated using Wilcoxon rank sum test with Bonferroni correction. *p<0.05 **p < 0.01 ***p < 0.001.

**Figure S5.**
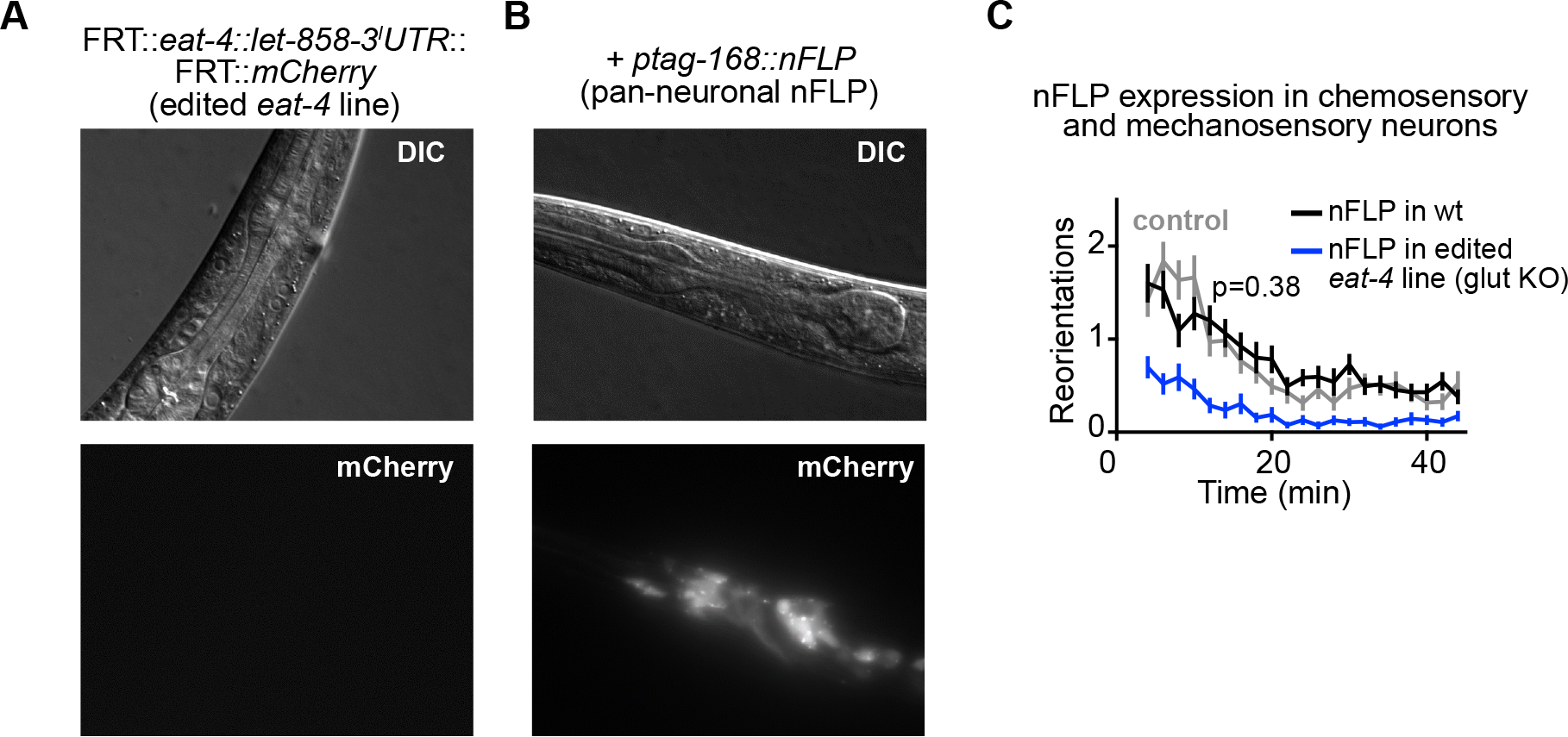
mCherry expression following *eat-4* excision with pan-neuronal nFLP, and nFLP expression control, Related to Figure 4. (A) Differential interference contrast (DIC) and mCherry fluorescence images of animals with edited endogenous *eat-4* (vGLUT1) locus. In the edited strain there is no mCherry expression, confirming that the *let-858*-3̍-UTR stops transcription. (B) DIC and mCherry images for edited *eat-4* strain following pan-neuronal nFLP expression (*ptag-168:::nFLp*). mCherry is broadly expressed throughout the nervous system. (C) Control experiment for flippase expression. Off-food foraging of wildtype animals expressing nFLP in chemosensory and mechanosensory neurons is indistinguishable from wildtype controls. This suggests that the local search defect following chemosensory and mechanosensory glutamate knockout is not caused by artifacts from flippase expression. (control, n=60; chemo+mechano nFLP in wt, n=63). p-values calculated using Wilcoxon rank sum test with Bonferroni correction.

**Figure S6.**
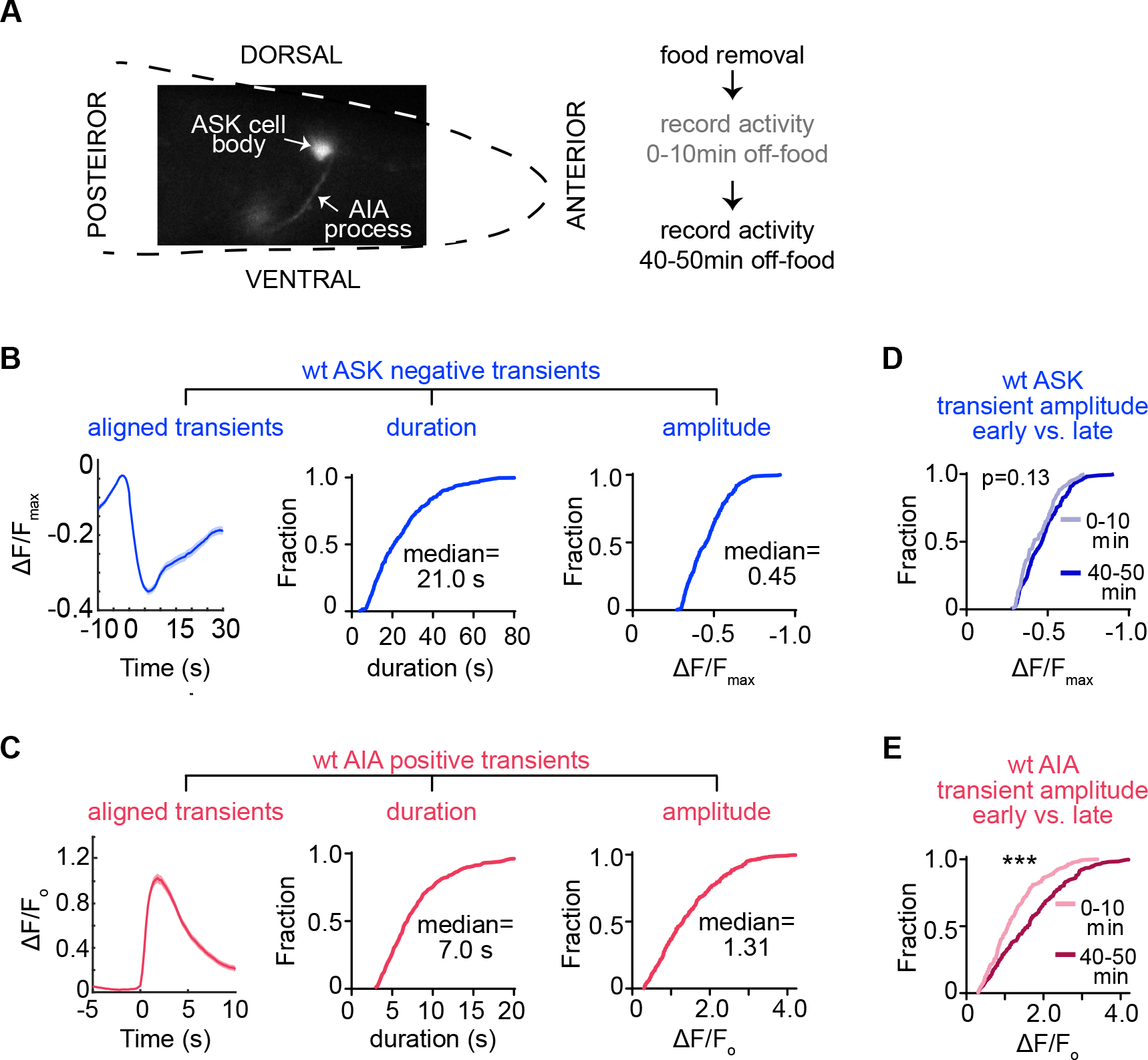
Calcium imaging and quantification, Related to Figure 5. (A) Experimental configuration for calcium imaging of ASK and AIA after food removal. GCaMP5A fluorescence levels were measured in the ASK cell body and in the AIA axon, either individually or simultaneously, 0-10 min after food removal and again 40-50 min after food removal. The animal is physically restrained within a custom-build microfluidic chamber that permits normal responses to chemosensory cues. Because the body is compressed in this environment, we did not attempt to record the activity of the mechanosensory circuit. (B) Average aligned traces, cumulative duration, and cumulative amplitude of ASK spontaneous negative transients in wildtype animals (n=18 animals, 327 transients). Data obtained from imaging ASK individually. (C) Average aligned traces, cumulative duration, and cumulative amplitude of AIA spontaneous positive transients in wildtype animals (n=21 animals, 465 transients). Data obtained from imaging AIA individually. (D) Cumulative ASK negative transient amplitude early (0-10 min, n=111) vs. late (40-50 min, n=216) after food removal for wildtype animals. p-value calculated using Kolmogorov-Smirnov test. Data obtained from imaging ASK individually. (E) Cumulative AIA positive transient amplitude early (0-10 min, n=210) vs. late (40-50 min, n=255) after food removal for wildtype animals. ***p<0.001 by Kolmogorov-Smirnov test. Data obtained from imaging AIA individually.

**Figure S7.**
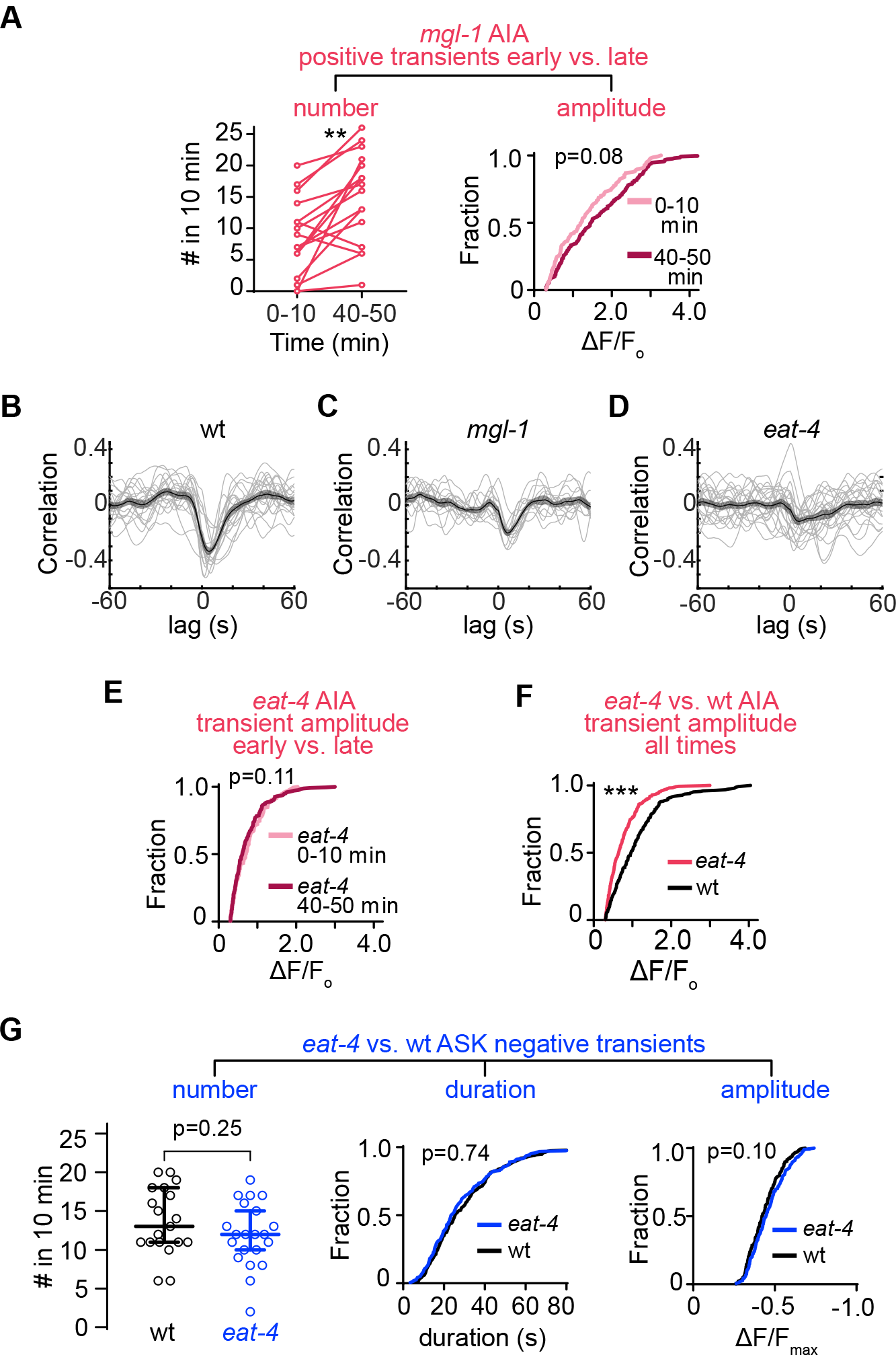
Calcium imaging in *mgl-1* and *eat-4* mutant animals, Related to Figure 5. (A) (Left panel) Number of AIA positive transients per animal early (0-10 min) vs. late (40-50 min) after food removal for *mgl-1* mutants (n=16). **p<0.01 by Wilcoxon signed rank test. (Right panel) Cumulative AIA positive transient amplitude early (0-10 min, n=137) vs. late (40-50 min, n=240) after food removal for *mgl-1* animals. p-value calculated using Kolmogorov-Smirnov test. Data obtained from imaging AIA individually. (B-D) Cross-correlation of ASK and corresponding lagged AIA traces 40-50 min after food removal in wildtype (B) (n=19), *mgl-1* mutants (C) (n=12) and *eat-4* (D) mutants (n=21). We selected 40-50 min because all animals show at least some spontaneous activity during this period. Data presented as mean ± s.e.m. Individual traces shown as thin lines. Data obtained from imaging ASK and AIA simultaneously. (E) Cumulative AIA positive transient amplitude early (0-10 min, n=179) vs. late (40-50 min, n=250) after food removal for *eat-4* mutants. p-value calculated using Kolmogorov-Smirnov test. Data obtained from imaging ASK and AIA simultaneously. (F) Cumulative AIA positive transient amplitude for wildtype (n=248 transients) vs. *eat-4* mutants (n= 421 transients). ***p<0.001 by Kolmogorov-Smirnov test. All data obtained from imaging ASK and AIA simultaneously. (G) Number (left), cumulative duration (middle), and cumulative amplitude (right) of ASK negative transients are similar in wildtype and *eat-4* mutants (data analyzed from 40-50 min period to match cross-correlations in panels B-D and aligned ASK transients in Fig. 5J). p-value on left panel calculated using Wilcoxon rank sum test. p-values on middle and right panels calculated using Kolmogorov-Smirnov test.

**Table S1.**
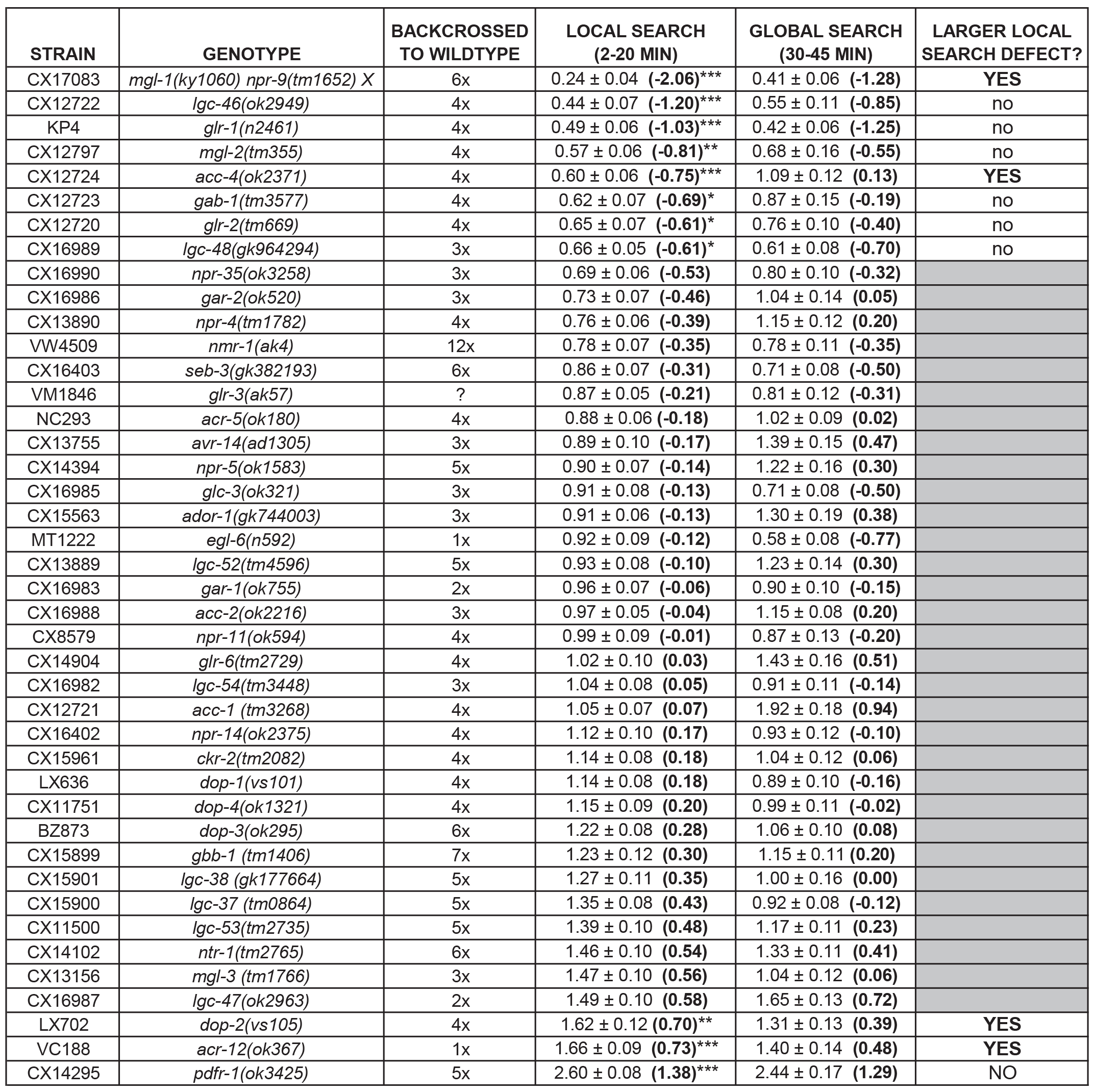
Results for candidate genetic screen. Strain name, genotype, number of times backcrossed to wildtype animals, and screen results. Since each mutant strain was matched to different control animals, we defined a ratio for each mutant strain by counting the number of reorientations of each mutant animal during the local search (2-20 min) or global search (30-45) time window and dividing it by the average number of reorientations of matched wildtype animals. Table shows mean ratio ± SEM for each mutant strain. In parentheses bolded is the log2 values for the ratio, which are plotted in Figure 2. To analyze the screen results, we found mutants that met two criteria: (1) the mutant was statistically different from wildtype during local search by at least 1.5-fold change (abs(log_2_(ratio)) > 0.58). Mutants that met this first criteria have their p-value result (asterisks) indicated in the ‘local search’ column, (2) local search was affected more than global search, which we assessed by comparing the two ratios. Strains meeting both criteria are indicated in the right column and marked with their local search p-value result (asterisks) in Figure 2. p-values calculated using Wilcoxon rank sum test with Bonferroni correction. * p<0.05 **p < 0.01 ***p < 0.001. Statistics were conducted by comparing the number of reorientations of mutant animals to matched controls during local search (see Methods).

**Table S2.**
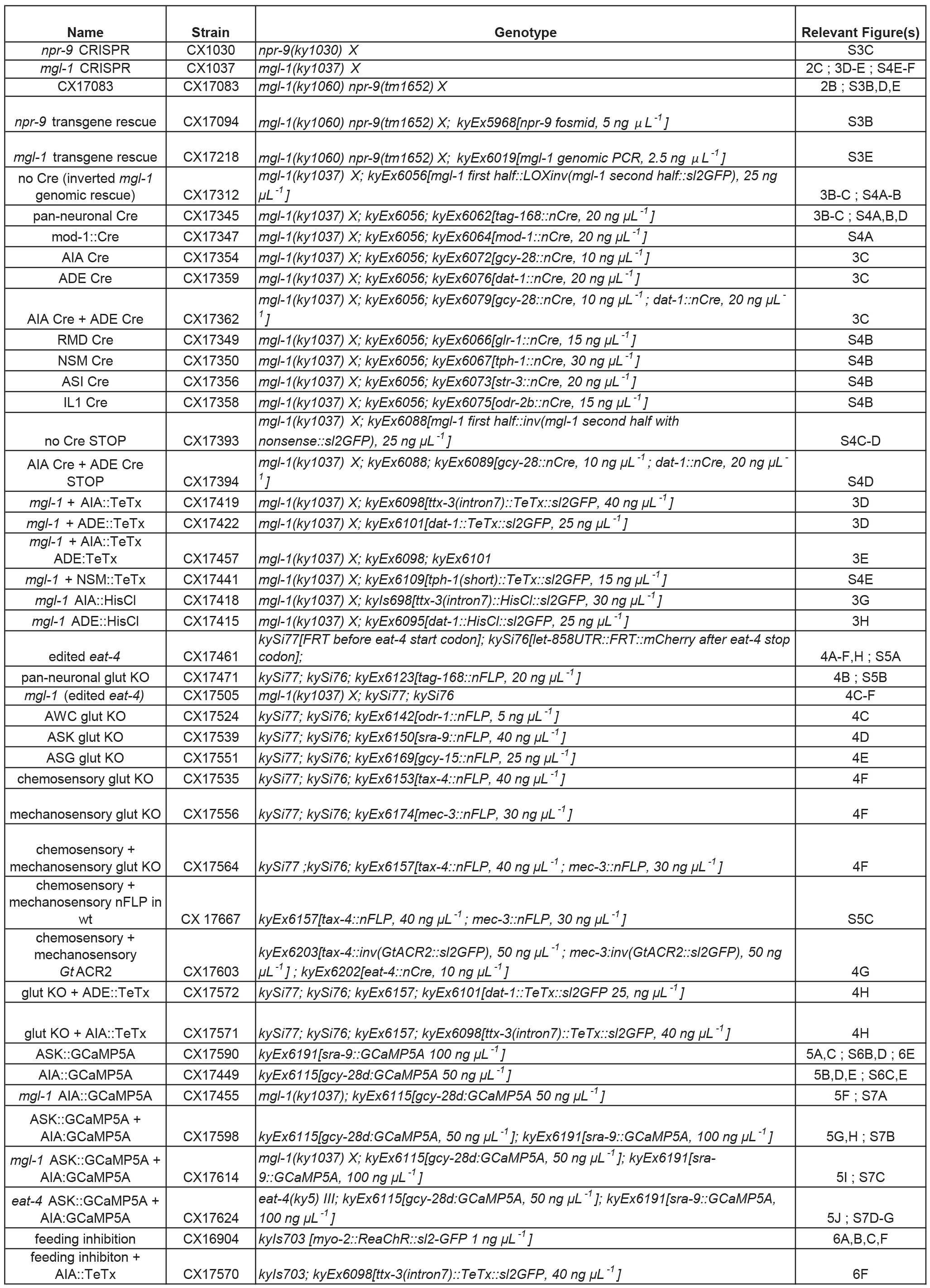
Strains generated for this study.

